# Anabolic Factors and Myokines Improve Differentiation of Human Embryonic Stem Cell Derived Skeletal Muscle Cells

**DOI:** 10.1101/2022.02.08.477036

**Authors:** Travis Ruan, Dylan Harney, Yen Chin Koay, Lipin Loo, Mark Larance, Leslie Caron

**Author notes:** <u>Correspondence</u>: Leslie Caron.

## Abstract

Skeletal muscle weakness is linked to many adverse health outcomes. Current research to identify new drugs has often been inconclusive due to lack of adequate cellular models. We have previously developed a scalable monolayer system to differentiate human embryonic stem cell (hESC) into mature skeletal muscle cells (SkMC) within 26 days without cell sorting or genetic manipulation. Here, building on our previous work, we show that differentiation and fusion of myotubes can be further enhanced using the anabolic factors testosterone (T) and follistatin (F) in combination with a cocktail of myokines (C). Importantly, combined TFC treatment significantly enhanced both hESC-SkMC fusion index and expression of various skeletal muscle markers including the motor protein Myosin Heavy Chain (MyHC). Transcriptomic and proteomic analysis revealed oxidative phosphorylation as the most up-regulated pathway and a significantly higher level of ATP and increased mitochondrial mass were also observed in TFC-treated hESC-SkMCs, suggesting enhanced energy metabolism is coupled to improved muscle differentiation. This cellular model will be a powerful tool for studying *in vitro* myogenesis and for drug discovery to further enhance muscle development or treat muscle diseases.

## Introduction

Skeletal muscle is the most abundant tissue in the human body, making up around 40% of total body weight. Skeletal muscle is essential for movement and metabolic health and diseases of muscle function can arise due to genetic mutation, metabolic or neuromuscular dysfunction, or natural aging[1,2]. Skeletal muscle disorders are linked to many adverse health outcomes such as impaired mobility, increased falls, fractures, frailty, diminished quality of life and premature death[3] and research to identify new drugs has often been inconclusive due to lack of adequate skeletal muscle models.

Due to species differences, animal models and rodent cell lines (e.g. C2C12 myoblasts, L6) do not accurately reflect all aspects of human muscle development[4,5]. Primary myoblasts obtained from patients’ biopsies have often been used for research but are limited in number, phenotypically diverse and have poor expandability, restricting their application. Human pluripotent stem cells (hPSC), on the other hand, offers a major advantage for studying human skeletal muscle development. Their capacity to proliferate indefinitely and differentiate into most cell types of the human body make them an excellent and renewable source of human skeletal muscle cells (SkMC). In recent years, hiPSC (human induced pluripotent stem cell) and hESC (human embryonic stem cell)-derived models from patients with muscular disease have become useful tools for modeling a large spectrum of inherited neuromuscular diseases[6,7,8]. While initially challenging and very inefficient, recent protocols utilize small molecules to recapitulate the embryonic development of skeletal muscle[6,7,9,10]. However, the majority of these protocols give rise to heterogeneous cell populations[11,12,7] and their reproducibility in multiple hPSC lines have proven a challenge[6,10]. We have previously developed a monolayer system to efficiently and reproducibly differentiate hPSCs into a pure and functional population of SkMC[6]. This protocol produced myosin heavy chain (MyHC) positive SkMC within 26 days that can be expanded in large quantities and has great potential for the study of human skeletal muscle development and diseases or drug screening. Our system has been efficiently reproduced in over 40 hPSC lines and is now used extensively in multiple laboratories [13–16]. However, myotubes remained thin and contained a small number of nuclei, suggesting that the last stage of the differentiation can be further improved.

To improve our skeletal muscle differentiation protocol, we investigated the effect of well-known anabolic compounds Testosterone (T), Follistatin (F) as well as a Cocktail of myokines (C) on hESC-SkMC. While these growth factors and hormone have been extensively studied *in vivo* or in primary SkMC, their combinatorial effect in hPSC differentiation into SkMC has not be reported. Here we show that a combination treatment of T, F and C (TFC) had a significant and synergistic effect on hESC-SkMC terminal differentiation and enhanced both hESC-SkMC fusion index and expression of various skeletal muscle markers. Transcriptomic and proteomic analysis revealed oxidative phosphorylation as the most up-regulated pathway, suggesting these cells also have greater capacity for energy metabolism.

## Materials and Methods

### Human Stem Cell Culture

GENEA002 and GENEA016 cell lines were obtained from Genea Biocells Ltd (Sydney, Australia). The ATCC-BXS0116 hiPSC line was used in this study (from ATCC, ACS1030). H9 cell line was obtained from WiCell institute.

GENEA016, GENEA002, H9 hESC and ATCC-hiPSC were cultured on Matrigel^®^ (Corning) coated plates in mTeSR™1 (StemCell Technologies) supplemented with 0.5% Penicillin-Streptomycin (ThermoFisher Scientific). Cells were grown in a 37°C incubator with 10% O_2_ and 5% CO_2_ and passaged every 3-4 days as needed.

### Skeletal Muscle Cell Differentiation

hPSC were differentiated into skeletal muscle cells following the protocol described in Caron et al[6]. Briefly, cells were seeded at a density of 2500 cells per cm^2^ onto Collagen Type I (Sigma-Aldrich) coated plates in Skeletal Muscle Induction Media (5% horse serum, 3μM CHIR99021, 2μM ALK5 inhibitor, 10ng/mL hr-EGF, 10μg/mL insulin, 0.4μg/mL dexamethasone, and 200μM ascorbic acid) for 10 days with media change every 2-3 days. At day 10, cells were dissociated and re-plated onto new Collagen Type I coated plates at 2500 cells per cm^2^ in Skeletal Muscle Myoblast Media (5% horse serum, 10μg/mL insulin, 10ng/mL hr-EGF, 20ng/mL hr-HGF, 10ng/mL hr-PDGF, 20ng/mL hr-bGFG, 20μg/mL oncostatin, 10ng/mL IGF-1, 2μM SB431542 and 200μM ascorbic acid) with media change every 2-3 days. At day 20, cells were switched to Skeletal Muscle Myotube Media (10μg/mL insulin, 20μg/mL oncostatin, 50nM necrosulfonamide, and 200μM ascorbic acid) with media change every 2-3days. The Skeletal Muscle Myotube Media is used for the culture and differentiation of non-treated myotubes (NTC). For TFC treatment, cells were cultured in Skeletal Muscle Myotube Media with the addition of 400ng/mL testosterone, 300ng/mL follistatin, 1mM creatine, 150ng/mL Il6, 20ng/mL Il4, 20ng/mL BDNF and 25ng/mL VEGF (Supp Table S2). For all myotubes differentiation experiments performed in this manuscript, cells were differentiated (with or without treatment) for 96 hours before analysis.

### Immunofluorescence Staining

Cells were washed with PBS and fixed with 4% formaldehyde for 15 minutes at room temperature followed by three times PBS wash and blocking in 1% BSA/PBS for 30 minutes. Cells were stained with primary antibody in permeabilisation buffer (1% BSA and 0.1% Triton-X 100 in PBS) and incubated overnight in 4C. On the next day, cells were washed three times with PBS and incubated with 1:500 Hoechst and appropriate secondary antibody in permeabilisation buffer for one hour at room temperature. The cells were washed with PBS three times before imaging.

### High Content Imaging and Analysis

Plates were imaged on Opera Phenix™ High-Content Screening System (Perkin Elmer) at 20x magnification and 49 fields (7 x 7) were imaged on every well. This imaging depth captures approximately 90% total area of an individual well. Image analysis was performed with Opera Phenix™’s analysis software Harmony using custom built myotube analysis pipeline. Briefly, MyHC were determined by MF20 signal and a thresholding filter was applied to remove non-specific and background signal. The dimension of MF20 is therefore considered as indicative of MyHC size. Similarly, nuclei were identified by Hoechst signal followed by a threshold filtering to remove background noise. Fusion index is calculated as the number of nuclei within MyHC+ area divided the number of total nuclei.

### Calcium Imaging

Calcium imaging protocol was adapted from[17]. Briefly cells were loaded with FURA-2AM (Invitrogen) in Hank’s Balanced Salt Solution (HBSS) for 30 minutes in the dark at 37C. Cells were washed three times with HBSS and maintained at 37C for 15 minutes prior to imaging on Nikon TI Live Cell Microscope (Nikon). Bath application of agonist (3mM nicotine) were performed after 70 seconds of baseline recording. The fluorescence ratio (F340:F380) were extracted from cells using Nikon NIS-Element software and presented as normalized mean ± SEM.

### ATP Determination Assay

ATP determination was performed using the ATP Determination Kit (A22066, Thermo Fisher) following the manufacturer’s protocol. Eight different batches of myoblasts were differentiated separately and subsequently transferred on individual wells of a new plate and ATP determination performed. Briefly, cells were lysed and supernatants incubated with recombinant firefly luciferase and its substrate D-luciferin. This assay is based on luciferase’s absolute requirement for ATP in producing light. Samples were read using a luminescence plate reader and ATP level was normalized to protein level using bicinchoninic acid protein assay.

### Mitotracker™ Deep Red Staining

Mitochondrial mass was determined using the Mitotracker™ Deep Red FM fluorescent dye (M22426, Thermo Fisher) following the manufacturer’s protocol. Briefly, cells were incubated with Mitotracker for 40 minutes in three concentrations (0/50/250nM) to allow permeabilization and imaged using Opera Phenix™. Images were analysed using Opera Phenix™’s analysis software Harmony using custom built Mitotracker analysis pipeline.

### RT-qPCR

Cells were collected from three different batches of D24 myotubes differentiated on different days. RNA were extracted using FavorPrep™ Blood/Cultured Cell Total RNA Kit (Fisher Biotec) and quantified using Qubit™ RNA BR Assay Kit (Thermo Scientific). cDNA was synthesized using iScript™ Reverse Transcription Supermix for RT-qPCR (Bio-Rad) and RT-qPCR was performed using SYBR™ Select Master Mix (Thermo Scientific) on LightCycler^**®**^Instrument II (Roche). Full list of primers for RT-qPCR are available on Table S3.

### RNASeq

All RNASeq experiment was performed in triplicate from three independent biological replicates. RNA were extracted and quantified as described above. RNA was treated with DNAse in solution using On-Column DNase I Digestion Set (Sigma-Aldrich) and maintained with Ribosafe RNAse Inhibitor (Bioline). Quality and quantity of RNA were assessed by Nanodrop (Thermo Scientific), Qubit (Thermo Scientific) and Bioanalyzer (Agilent). Library preparation and sequencing were performed by Novogene. STAR aligner[18] was used for mapping sequence reads to the human genome (hg38 assembly), allowing up to three mismatches and retaining only reads that align to unique locations. Ensembl gene models[19] was used for quantifying gene expression from mapped reads using featureCounts[20] and genes that are lowly expressed (less than two samples with counts >10) were removed from subsequent analysis. Raw read counts were analysed in RStudio using DESeq2 and differential expression was assessed[21]. Genes with an adjusted p-value of <0.05 (Benjamani Hochberg corrected) were assessed as differentially expressed.

### Proteomics

All proteomics experiment was performed in triplicate from three independent biological replicates. Proteomics protocol was adapted from[17]. Cells were lysed in 4% sodium deoxycholate (SDC) buffer and heat inactivated for 10 minutes at 100C. Sonicated samples were quantified using bicinchoninic acid protein assay (Thermo Scientific) following the manufacturer’s instructions to determine protein concentration. Tandem mass spectrometry was carried out on a Q-Exactive Mass Spectrometer (Thermo Scientific). Raw data were processed using MaxQuant using human UniProt database. The quantification of proteins in LFQ intensity values were log-transformed (base 2). We next filtered proteins requiring all biological replicates to be quantified in at least one condition. After filtering, the missing values were then imputed using the tail imputation method with a Gaussian distribution of 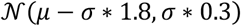 as in [22]. The imputed data were next converted to ratios relative to the control condition and normalized using Combat[23] for removing additional unwanted variation. Differentially regulated proteins were determined using ANOVA test with an adjusted p-value of <0.05.

### Metabolomics

All metabolomics experiment was performed in triplicate from three independent biological replicates. Cells were harvested and subjected to metabolite extraction as described in[24], with some minor modifications. Cells were washed twice with cold sodium chloride and scraped in 500uL 50% (vol/vol) methanol:water mixture containing internal standards of 10mM phenylalanine-d_8_, valine-d_8_ and thymine-d_4_ on ice (4C). 500uL of chloroform was added to the extracts and vortexed. The aqueous phase was separated from the insoluble and organic layers by centrifugation at 16,000g, 4C for 20 minutes. The upper aqueous phase was subjected to drying using SpeedVac Vacuum concentrator and resuspended in acetonitrile:methanol:formic acid (75:25:0.2; v/v/v, HPLC grade; Thermo Fisher Scientific) for HILIC-MS method and in acetonitrile:methanol (75:25; v/v HPLC grade; Thermo Fisher Scientific) for AMIDE-MS method. Every extraction condition was prepared in three biological replicates. Samples were analysed by liquid chromatography-tandem mass spectrometry (LC-MS/MS).

LC-MS/MS analysis was performed using an Agilent Infinity 1260 LC coupled to an AB Sciex QTRAP 5500MS. LC separation for AMIDE-MS method was achieved on a XBridge Amide column (4.6mm x 100mm, 3.5uml Waters Australia) at ambient temperature using buffer A (95:5; v/v) water:acetonitrile containing ammonium hydroxide and ammonium acetate both at 20mM (pH 9.3) and buffer B (100% acetonitrile). LC separation for HILIC-MS method was performed on Atlantis^®^ HILIC column (2.1mm x 150mm, 3um; Waters Australia) using buffer A (water containing formic acid (0.1%) and ammonium formate (10mM)) and buffer B (acetonitrile containing formic acid (0.1%)).

All raw data files (Analyst software, version 1.6.2; AB Sciex, Foster City, CA, USA) were acquired and imported into Multi-Quant™ 3.0 Software for MRM Q1/Q3 peak integration.

### Pathway Analysis

Pathway analysis of transcriptomic and proteomic data were performed on Ingenuity Pathway Analysis (IPA) using genes with an adjusted p-value of <0.05 as input. Gene Set Enrichment Analysis (GSEA) was used to determine whether there were significant differences between treatments using default parameters.

### Availability of Data and Material

The RNASeq data were deposited into the National Center for Biotechnology Information (NCBI) Gene Expression Omnibus (GEO) database with accession number GSE174144.

Raw proteomics data have been deposited to the ProteomeXchange Consortium (http://proteomecentral.proteomexchange.org) via the PRIDE partner repository with the dataset identifier PXD025906, username, password: fCoIAiDN. The datasets generated during and/or analysed during the current study are available from the corresponding author on reasonable request.

### Ethics Approval

Not applicable

## Results

### hESC Differentiate into Functional SkMC *in vitro*

We first differentiated GENEA016 hESC into hESC-SkMC following the protocol described by *Caron et al* [6] and confirmed the myogenic identity of these cells by immuno-staining for markers representative of each stage (Fig. 1A-B and Supp. 1A-C). During myogenic lineage induction (Day 1 – 10), cells stained positive for PAX3 and PAX7, two transcription factors known for their role in early phase of myogenesis [25] (Supp. 1A). After the second phase of differentiation (Day 10 – 20), 58.2% of the cells stained for MyoD1 at D20, the master regulator of myogenesis indicative of myogenic lineage commitment [26] (Supp. 1B). After myoblasts elongation and fusion during the terminal phase of differentiation, 79.4% of myotubes stained positive for the skeletal muscle marker Myogenin (Supp. 1C) and myotubes expressed the sarcomeric proteins Dystrophin, alpha-Actinin and MyHC (Fig. 1B). As previously shown, MyHC (MF20 staining) is detected in 70% of differentiated hESC-SkMC (D24) and not in pre-differentiation hESC (D0) (Supp. 1D). hESC-SkMC expressed MYH3 (embryonic) and MYH8 (perinatal) but not MYH1 or MYH2 (adult) isoforms (Supp. 1D), indicating a relatively immature phenotype^6^.

**Figure. 1.**
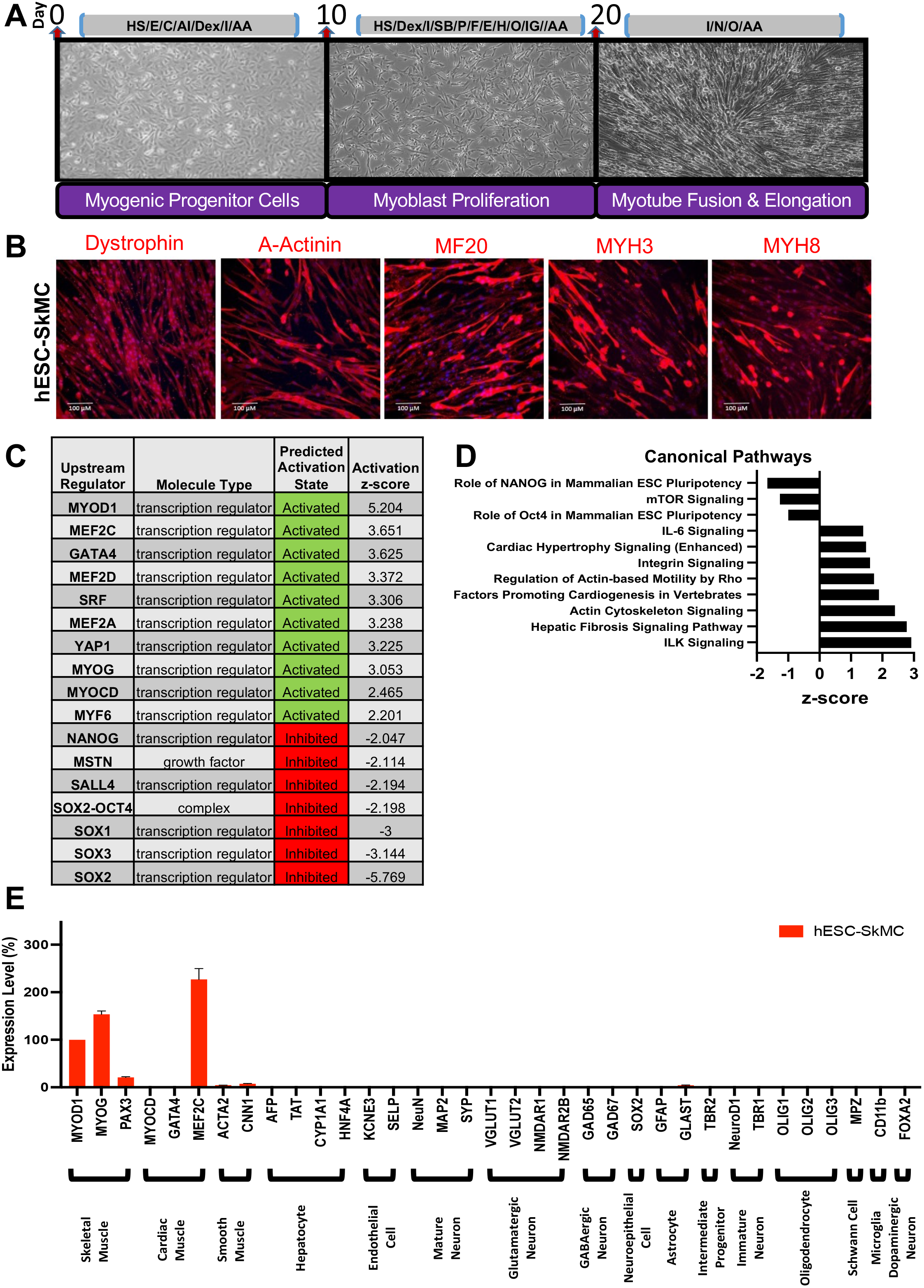
Skeletal muscle differentiation of hESC. (A) Differentiation protocol for the derivation of SkMC from hESC. HS = Horse Serum, E = hr-EGF, C = CHIR99021, AI = ALK5 Inhibitor, Dex = Dexamethasone, AA = Ascorbic Acid, I = Insulin, SB = SB431542, P = hr-PDGF, F = hr-FGFB, H = hr-HGF, O = Oncostatin, IG = hr-IGF1, N = Necrosulfonamide. (B) hESC-SkMC express high level of SkMC markers Dystrophin, a-Actinin, MF20 (MYH all isoforms), embryonic (MYH3) and perinatal (MYH8) myosin. (C) IPA transcriptomic analysis of upstream regulators between hESC and hESC-SkMC and their activation status. (D) IPA transcriptomic analysis of differentially regulated canonical pathways between hESC and hESC-SkMC. (E) RNASeq data showing hESC-SkMC express markers specific to skeletal muscle lineage. Shown is data pooled from 3 independent biological replicates.

We compared the transcriptomic and proteomic profile between hESC and hESC-SkMC and demonstrate appropriate expression pattern associated with cell differentiation including down-regulation of key pluripotency genes NANOG, POU5F1 and SOX2, and down-regulation of proliferation and cell cycle genes including MKI67 and CDK1 at the protein level (Supp. 1F). CDK4 was up-regulated as it has been shown to allow myogenic cells to recapture growth property without compromising differentiation potential[27]. Additionally, we performed IPA (Log2FC +/−4, padj<0.05) to identify the cascade of upstream transcriptional regulators that can explain the observed changes in gene expression. We assessed the expression level of numerous key muscle and pluripotency transcriptional factors, and their activation z-score suggest an enhancement of muscle differentiation pathways and inhibition of pluripotency pathways consistent with *in vitro* muscle differentiation (Fig. 1C). The core pluripotency network including OCT4, NANOG and SOX2 were predicted as inhibited. Similarly, Myostatin, a well-known myokine that inhibits myogenesis[28] was predicted as inhibited. In contrast, myogenesis transcription regulators, including all members of the myogenic regulatory factors (MRF) MyoD1, MYOG, MYF5 and MYF6 (also known as MRF4) were activated (Fig. 1C, Supp. File 1). These myogenic factors are involved in cell specification of the skeletal muscle lineage and are important in the generation of both developing and mature skeletal muscle[29]. These transcriptomic changes are associated with modulation of several signaling pathways (Supp. File 2). Some examples include down-regulation of pathways involved in cell pluripotency and proliferation, as well as up-regulation of those that functions in myocyte contractility (ILK Signaling)[30], cell-cell adhesion (Integrin Signaling)[31] and sarcomere integrity (Cardiac Hypertrophy Signaling)[32] (Fig. 1D). To confirm the cell purity of our differentiation system, we evaluated the expression of specific markers of various cell types by RNAseq in our differentiated population relative to MyoD1 expression level. Non-muscle markers were not expressed in hESC-SkMC. This includes markers for several neuronal lineages as well as astrocytes, oligodendrocytes, hepatocytes and endothelial cells (Fig. 1E). Cardiac and smooth muscle markers were also undetected in our hESC-SkMC, with the exception of MEF2C and CNN1. While MEF2C is a cardiac lineage marker, it is also expressed in skeletal muscle during development[8], which explains its presence in our hESC-SkMCs. CNN1 is highly expressed in smooth muscle but can also be detected in skeletal muscle[33] (Supp. 1G). All together, these results demonstrate the specificity of our skeletal muscle differentiation method and confirm what we and others previously reported [6,13]. As previously described by *Caron* et al., 2016, the remaining cells that do not stain positive for MF20 in the myotube culture at D24 are unfused myogenic progenitors or myoblasts[6,34]. Lastly, to assess whether hESC-SkMC were functionally responsive *in vitro*, we stimulated the cells with nicotine (3mM) and performed calcium imaging. hESC-SkMC were able to respond and we observed calcium transients upon nicotine stimulation (Supp. 1H).

### MyHC Expression in hESC-SkMC is Enhanced by Testosterone, Follistatin, Cocktail of myokines and the Combination Treatment

To improve the final stage of our differentiation protocol, we selected factors with reported positive effects on skeletal muscle growth or strength. These include creatine, a non-protein nitrogenous compound known to increase skeletal muscle strength and performance[35,36], and the myokines IL-4, IL-6 (interleukin 4, -6), BDNF (Brain-Derived Neurotrophic Factor) and VEGF (Vascular Endothelial Growth Factor), which are up-regulated by exercise[37,38] and regulate muscle mass and function[39]. For each of these factors, we chose the concentration that is commonly used in the literature. We noticed slight changes in myotubes morphology when these five compounds were added together (data not shown). We therefore used these compounds as a mixture that we refer to as “Cocktail of myokines” (C) in this study. Additionally, based on their strong anabolic effects, we also selected the well-known steroid hormone testosterone (T) and the myostatin inhibitor follistatin (F)[40] as potential muscle differentiation enhancers. We assessed the effect of T, F, C and the combined treatments (TFC) on hESC-SkMC. Brightfield image showed average myotube size was greater following treatment indicating improved terminal differentiation and fusion capacity (Supp. 1I). We next performed a detailed morphological analysis of treated and non-treated hESC-SkMC. Since myosin is the most abundant protein in the muscle and the expression level of this protein reflects skeletal muscle size and strength[41] we therefore investigated myosin abundance in hESC-SkMC following each treatment. MyHC expression was detected by immunofluorescence using MF20 antibody (recognizing all MyHC isoforms) in hESC-SkMC 96 hours after treatment with T, F, C, TFC or vehicle control. Size of MyHC+ areas were analyzed using the Perkin Elmer Opera Phenix™ automated high-content screening system that produces high-throughput imaging and image quantification. Briefly, input images were split into individual channels and a threshold filter was applied to remove background noise. Signals that passed the threshold were subsequently quantified (Fig. 2A). While individual treatments had no significant effect on MyHC+ areas when compared to non-treated cells (NTC) (Fig. 2B-C), MyHC+ area was significantly increased in TFC-treated cells (Fig. 2C). MyHC+ myotubes were also thicker in TFC-treated cultures compared to NTC (Fig. 2B-C). As myotubes are multinucleated, we quantified the number of nuclei within MyHC+ fiber and observed a significant increase in cells treated with TFC (Fig. 2E), associated with an increase of the total number of nuclei (Fig. 2D). In addition, we observed a significant increase in myotubes fusion index (calculated as the number of nuclei within MyHC+ area divided by the number of total nuclei) after treatment with TFC (Fig. 2F), demonstrating these cells have better fusion capacity. To assess consistency of this treatment we also evaluated TFC treatment on three additional hPSC lines. We observed similar results in a hiPSC line (ATCC-hiPSC) and a hESC line (GENEA002) and a similar trend in one other hESC lines (H9 (WA09)) (Supp. 2A-C). RNAseq analysis also revealed that TFC treatment did not affect the purity of our hESC-SkMC (Supp. 2D).

**Figure. 2.**
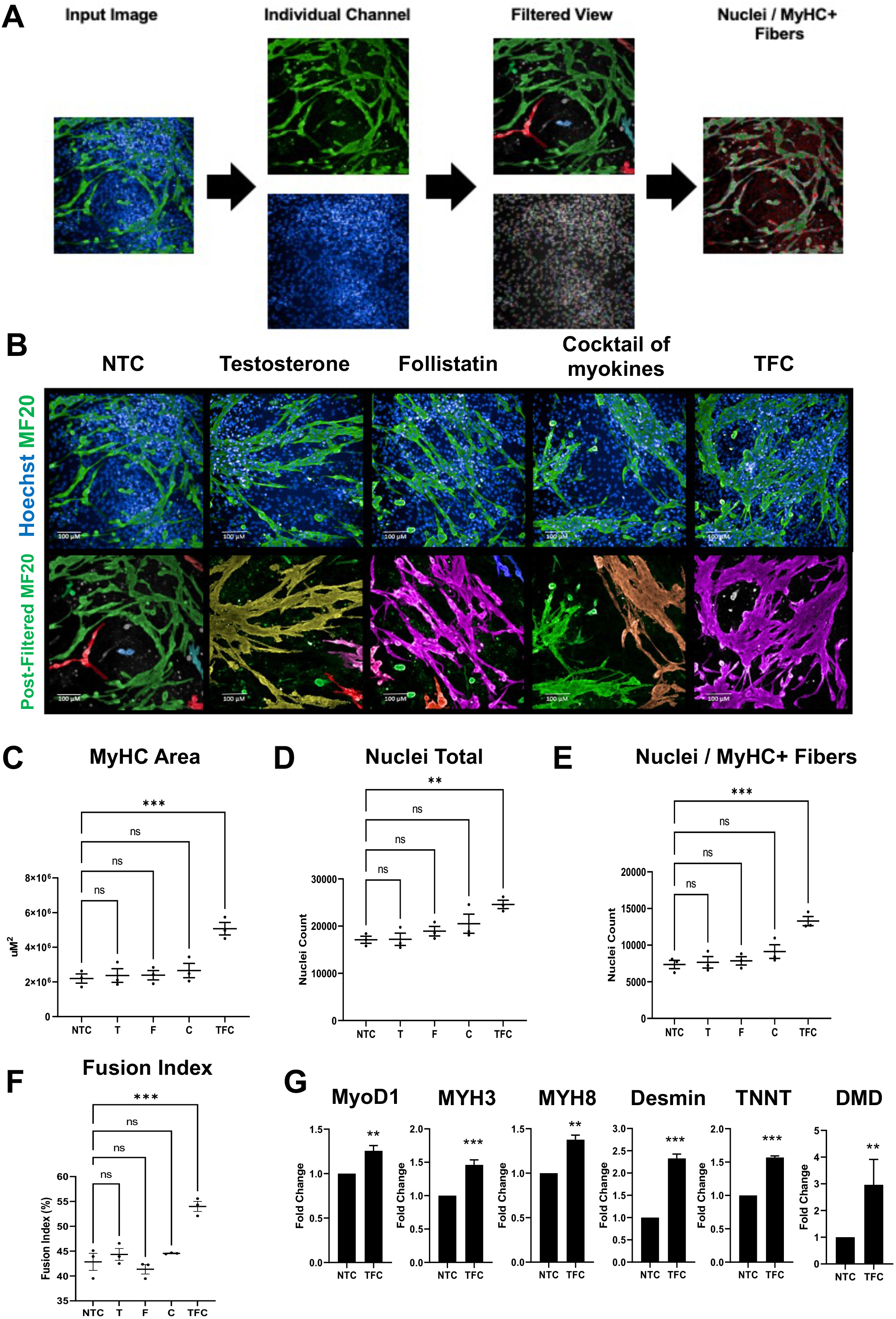
Anabolic factors and myokines enhance terminal differentiation of hESC-SkMC. (A) Image analysis pipeline. (B) Example pre and post-filtered image of untreated and treated hESC-SkMC. (C-F) Image quantification of hESC-SkMC between different treatments for MyHC area (C), Nuclei total (D), Nuclei within MyHC+ fibers (E) and Fusion index (F). N = 3 for each condition, obtained as the average of four independent technical replicates over three independent experiments. Statistical analysis performed using One-way ANOVA with Benjamini-Hochberg FDR correction. *P < 0.05, **P < 0.01, ***P < 0.001. (G) TFC enhanced expression of several key myogenesis markers assessed by RT-qPCR. Statistical analysis performed using two tailed t-test. *P < 0.05, **P < 0.01, ***P< 0.001.

To evaluate the general effect of TFC on hESC-SkMC differentiation, we analyzed the expression of various skeletal muscle specific genes by RT-qPCR and found TFC enhanced the expression of MyoD1 (Myoblast Determination Protein 1), MYH3 (embryonic) and MYH8 (perinatal) myosin heavy chain isoforms, Desmin (muscle specific intermediate filament), and the sarcomeric structural proteins TNNT (Troponin T) and DMD (Dystrophin) (Fig. 2G). We also compared their responsiveness to nicotine stimulation (Supp. 3A). Although no difference was observed in the mean amplitude of calcium transient between NTC and TFC treated hESC-SkMC (Supp. 3B), TFC treated myotubes were more responsive to nicotine stimulation compared to NTC both in cell number and total surface area (Supp. 3C-D). Lastly, TGFβ signaling pathway has been shown to be an important regulator of myoblasts differentiation and its inhibition was reported to enhance skeletal muscle fusion efficiency in both primary SkMC and hPSC-SkMC [42,43]. We therefore compared the effect of TFC with the highly selective TGFβ inhibitor ITD-1 on hESC-SkMCs differentiation. In our system, we observed no significant increase of MyHC+ area and fusion index in cells treated with ITD-1, while TFC led to a significant increase of both MyHC+ area and fusion index (Supp. 3E-F). Collectively, these results demonstrate addition of TFC can further enhance terminal differentiation by promoting fusion and MyHC expression of hESC-SkMC.

### Skeletal Muscle Genes Expression in hESC-SkMC is Slightly Enhanced by TFC Treatment

To determine the gene expression change behind enhanced MyHC expression and skeletal muscle fusion, we performed RNASeq to assess the transcriptomic profile between NTC and treated hESC-SkMC (T/F/C/TFC). Three biological replicates corresponding to hESC-SkMC derived from three independent differentiation experiments were used for each of the treatments. Using padj <0.05 and Log2FC>1 as threshold, we observed 671 differentially expressed genes (DEG) in TFC treatment including 471 up-regulated and 200 down-regulated genes compared to NTC (Fig. 3A, Supp. File 3). Surprisingly, many of these DEGs observed in TFC were different from those in individual treatments (T, F or C) (Supp. 3G-H, Supp. File 4). To validate this transcriptomic profile, we performed RT-qPCR and confirmed the up-regulation of several top DEGs in TFC-treated hESC-SkMC that we chose based on their reported role in muscle development and function (Fig. 3B). These genes are involved in a variety of biological processes known to be associated with skeletal muscle functioning including metabolism of lipids (ALOX15, FABP4), interaction with vitamins (RBP4, VDR) and calcium binding (SCGN, DUOX2). Since TFC-treated hESC-SkMC have overall higher MyHC protein expression and increased expression of MYH3 and MYH8 (Fig. 2B, Fig. 2G), we next compared the expression of a number of myosin or sarcomere genes between NTC and TFC treated hESC-SkMC. Surprisingly, we did not observe any major differences (Fig. 3C). Moreover, TFC-treated cells have minimal levels of MYH1 and MYH2 expression, suggesting treatment does not promote maturation towards adult MyHC (Supp. 3I).

**Figure. 3.**
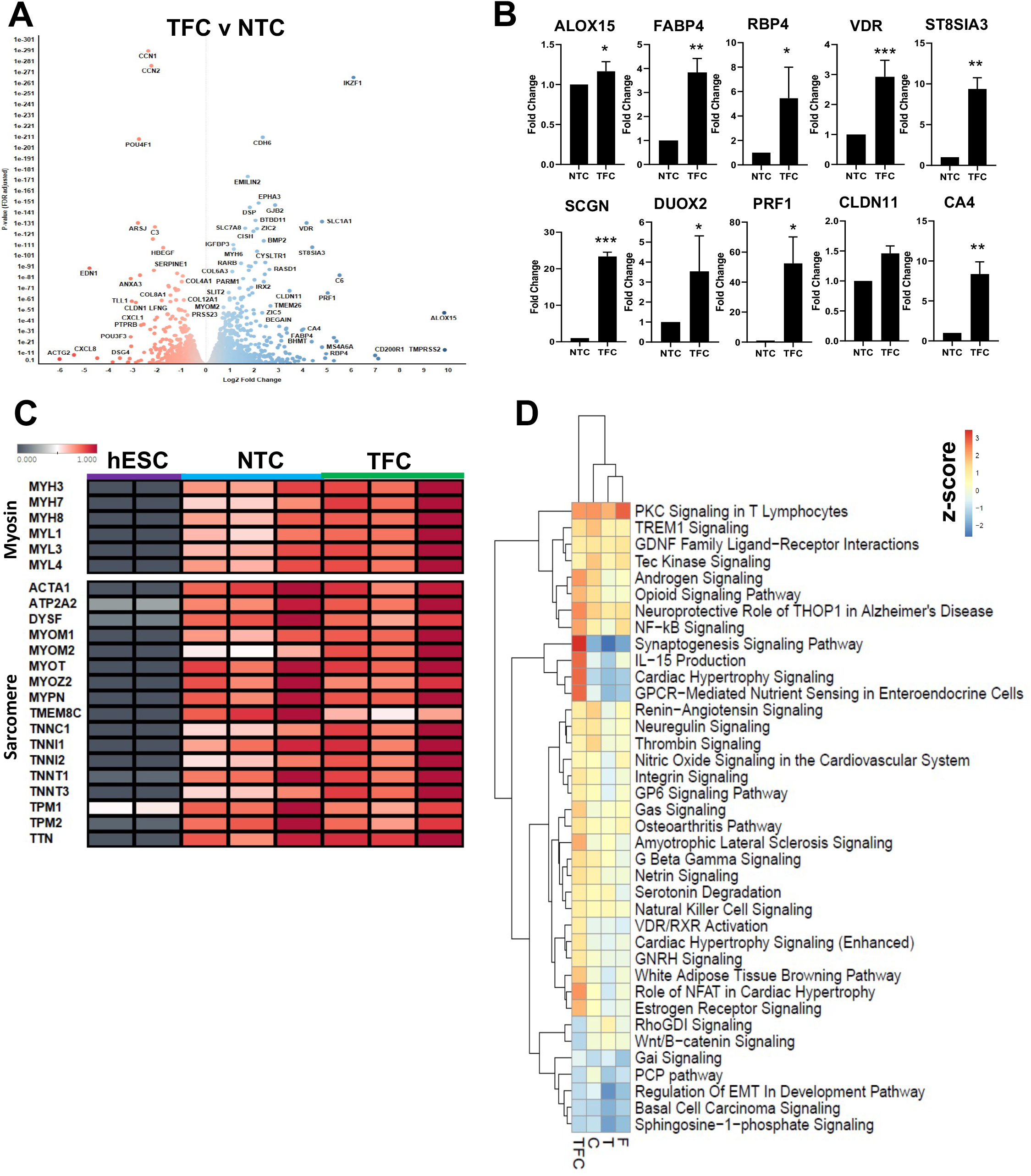
Transcriptomic profiling of NTC and TFC treated hESC-SkMC. (A) Volcano plot of differentially expressed genes between TFC and NTC. (B) RT-qPCR validation of several top differentially expressed genes as identified via RNASeq. Statistical analysis performed using two tailed t-test. *P < 0.05, **P < 0.01, ***P< 0.001. (C) Heatmap comparison of various myosin and sarcomere genes between hESC, NTC and TFC-treated hESC-SkMC. (D) IPA comparative analysis of differentially regulated pathways between treated hESC-SkMC compared to NTC. Shown is data pooled from three independent biological replicates.

### Differentially Regulated Pathways Show Common and Divergent Patterns

Since transcriptional expression of skeletal muscle genes (Fig. 3C) was not markedly enhanced by TFC treatment, we decided to identify the molecular pathways that were induced by TFC treatment. We compared the expression profile between treated hESC-SkMC (T/F/C/TFC) to NTC using IPA (Log2FC>1, padj<0.05) (Fig. 3D, Supp. File 5). DEGs were grouped into 222 pathways and we observed pathways that shared similar expression trends in all treatments such as PKC Signaling and Androgen Signaling. Interestingly, TFC also up-regulated several pathways that were not up-regulated by individual treatment, such as Cardiac Hypertrophy Signaling. Cardiac hypertrophy signaling is involved in sarcomere formation and includes the well-known myocyte enhancer MEF2C, which plays an important role during myogenesis[44].

### TFC Highly Enhanced Myosin and Sarcomere Gene Expression at the Protein Level

Myogenesis is closely associated with a high level of post-transcriptional modification events[45,46,47]. Since we did not observe major changes in skeletal muscle specific genes on the transcript level, we considered differential expression between NTC and TFC might be more noticeable at the protein level. We therefore performed proteomics and compared NTC and treated hESC-SkMC (T/F/C/TFC). We identified a large number of proteins (T = 195, F = 260, C = 214) that were not expressed in NTC but expressed upon individual compound treatment (Supp. 4A, Supp. File 6). Interestingly, a subset of 80 proteins was up-regulated in all three individual treatments. Gene Ontology pathway analysis showed these 80 proteins are involved in mitochondrial translation (57.14%) and Complex I Biogenesis (42.86%) (Supp. 4B), suggesting these individual treatments enhance cellular respiration rate and ATP production. Surprisingly, 1033 proteins were not expressed in NTC but present in TFC-treated SkMC (Supp. File 7). Among these, 803 proteins are only detected in TFC treatment and not in individual treatment (Supp. 4C-D). Gene Ontology analysis revealed these 803 proteins to be involved in diverse cellular activities including glutamine and L-alanine transport, multivesicular body organization, organelle biogenesis and maintenance and metabolism (Supp. 4E). Next, we compared the proteomic profile between TFC and NTC treated hESC-SkMC (Fig. 4A) and assessed differentially regulated pathways via GSEA (Supp. 4F-G, Supp. File 8). We observed highly up-regulated protein set clusters closely associated with muscle contraction, skeletal muscle development, and respiratory chain electron transport. In contrast, major down-regulated gene set clusters include condensed chromosome and regulation of mRNA processing, suggesting TFC treated hESC-SkMC might be less transcriptionally active than NTC.

**Figure. 4.**
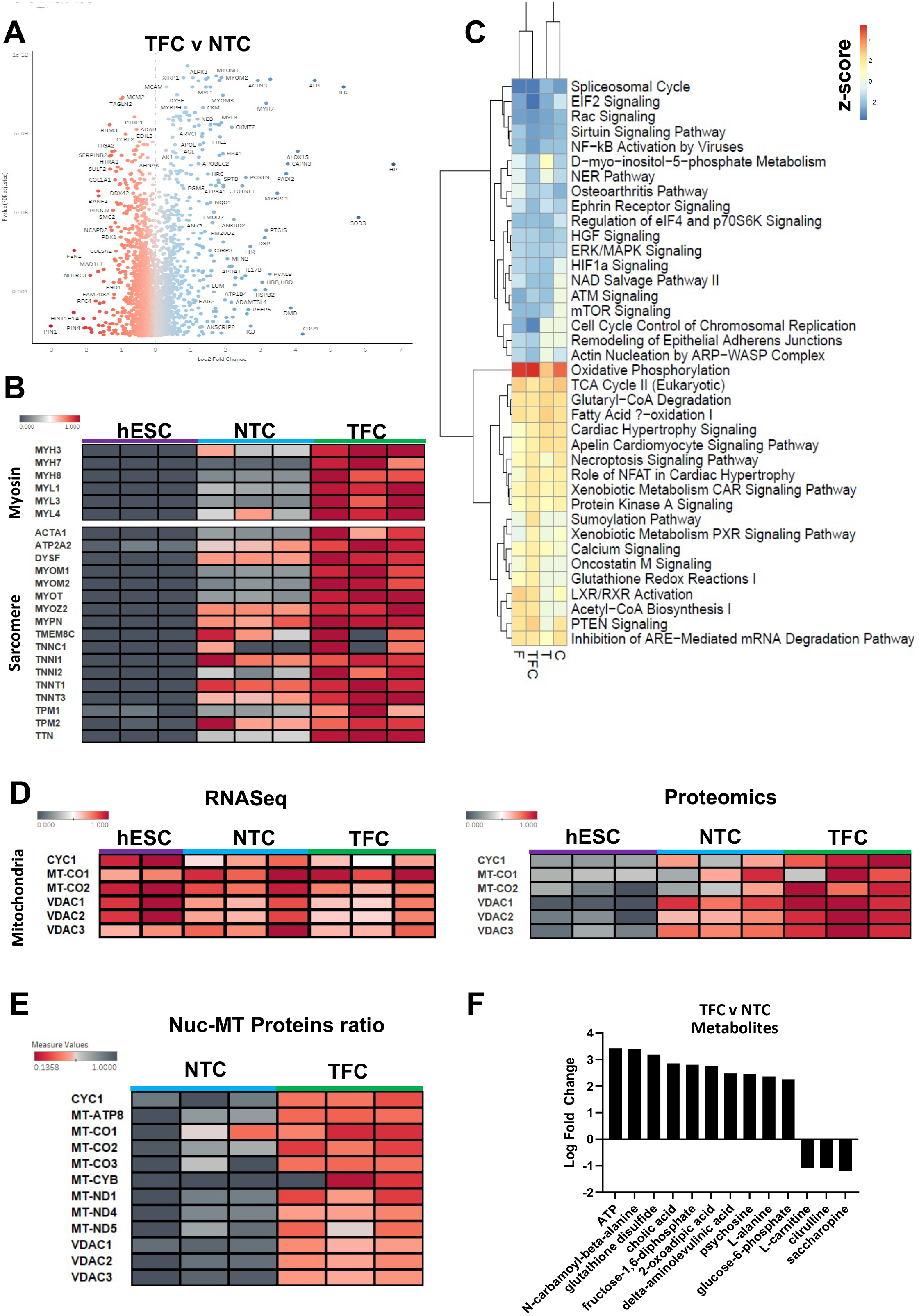
Proteomics profiling of NTC and TFC treated hESC-SkMC. (A) Volcano plot of differentially expressed proteins between TFC and NTC. (B) Heatmap comparison of various myosin and sarcomere proteins between hESC, NTC and TFC-treated hESC-SkMC. (C) IPA comparative analysis of differentially regulated pathways between treated hESC-SkMC compared to NTC. (D) Heatmap comparison of various mitochondrial genes at RNA (left panel) and proteins (right panel) level between hESC, NTC and TFC-treated hESC-SkMC. (E) Heatmap comparison of Nuclear (Histone H4) vs Mitochondrial proteins ratio in NTC and TFC-treated SkMC. (F) IPA metabolomics analysis (Log2FC>1, padj<0.05) of differentially expressed metabolites between TFC and NTC. Shown is data pooled from three independent biological replicates.

To determine whether the proteomic expression follows the transcriptomic profile, we evaluated the expression of skeletal muscle markers shown in Fig. 3C. Interestingly, while transcriptomic data showed no significant difference in skeletal muscle markers expression between NTC and TFC, we observed a significant change in these markers at the protein level. The expression of various myosin heavy and light chain isoforms (MYH3, MYH7, MYH8, MYL1, MYL3 and MYL4) and sarcomere structural proteins such as Titin (TTN) and Actin Alpha 1 (ACTA1) were significantly up-regulated in TFC compared to NTC (Fig. 4B). However, MYH1 and MYH2 (adult MyHC isoforms) proteins remain undetected after TFC treatment (Supp. 3I). Pluripotency (NANOG, POU5F1 and SOX2) and proliferative markers (MKI67) were not detected on the protein level in either NTC or TFC treated cells, indicating both protocols lead to a post-mitotic cellular state (Supp. 5A).

### Oxidative Phosphorylation is the Most Up-Regulated Pathway in Treated hESC-SkMC

To identify the molecular pathways associated with protein changes, we performed IPA (Log2FC>1, padj<0.05) comparison of expression profile between treated hESC-SkMC (T/F/C/TFC) to NTC and observed oxidative phosphorylation as the most up-regulated pathway in all treatments, suggesting these anabolic factors and myokines had a positive effect on enhancing cellular respiration in hESC-SkMC (Fig. 4C). In addition to oxidative phosphorylation, several other pathways known to be involved in energy metabolism, including TCA Cycle, Fatty Acid B-oxidation and Acetyl-CoA Biosynthesis were also up-regulated in all treatments compared to NTC (Fig. 4C). It is known that hESC primarily utilizes glycolysis for energy production and switches to oxidative phosphorylation as they differentiate into specialized cell type[48]. We did not observe significant difference in key mitochondrial genes expression when comparing NTC and TFC-treated hESC-SkMC by RNAseq (Fig. 4D). However, mitochondrial proteins were highly enriched in hESC-SkMC compared to hESC. Notably, mitochondrial protein expression level is much higher in TFC treated hESC-SkMC than untreated cells, suggesting that TFC promotes greater metabolic activity (Fig. 4D). We further assessed the expression of a large number of mitochondria related genes between conditions and observed a noticeable increased expression in TFC-treated cells at the protein but not mRNA level (Supp. 6A). Furthermore, we performed nuclear to mitochondrial protein ratio comparison by normalizing it to Histone H4 protein expression level. Normalised expression ratio demonstrate treated hESC-SkMC also showed higher mitochondrial proteins ratio compared to NTC, and the overall effect was greatest in TFC (Fig. 4E and Supp. 6B). We next performed metabolomics to assess the metabolites produced by NTC and TFC. Differential analysis revealed ATP as the most enriched metabolite in TFC treated hESC-SkMC compared to NTC, matching with the proteomics data, and is accompanied with up-regulation of metabolic pathways such as Glycolysis, Fatty Acid Activation and FAT10 Signaling Pathway (IPA Log2FC>1, padj<0.05) (Fig. 4F and Supp. 5B).

### TFC Treatment Enhanced Oxidative Phosphorylation in hESC-SkMC

To confirm TFC treatment can enhance oxidative phosphorylation in hESC-SkMC, we performed quantitative ATP determination (Fig. 5A). TFC treated hESC-SkMC has significantly higher level of ATP compared to NTC (1.47 fold change). We next assessed mitochondrial mass in NTC and TFC treated hESC-SkMC using the fluorescent dye Mitotracker™ Deep Red (250 nM) (Fig. 5B-F). In skeletal muscle, mitochondria form a very dense network making the visualization, count and assessment of individual mitochondria difficult.

**Figure. 5.**
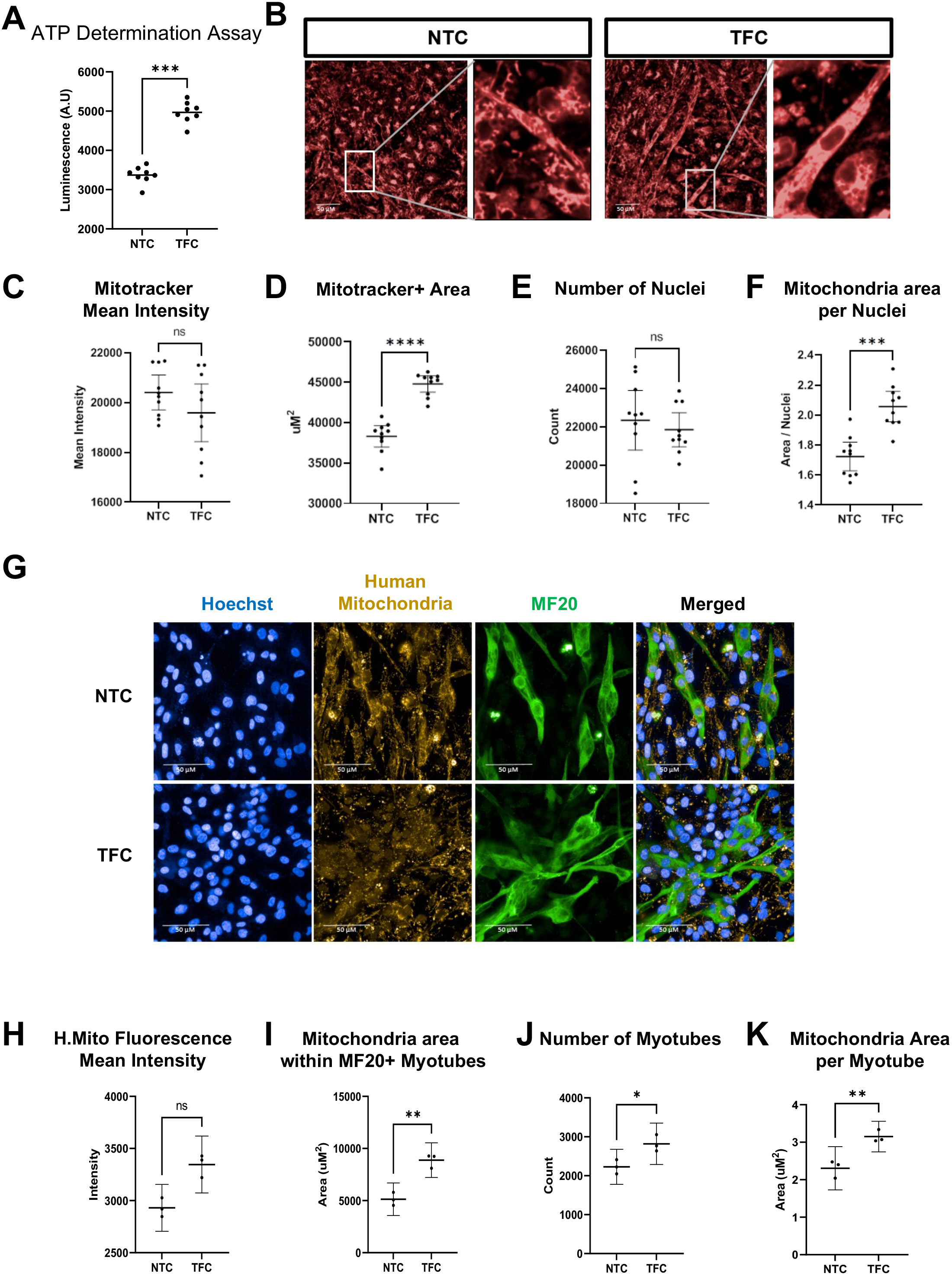
TFC treatment enhanced oxidative phosphorylation in hESC-SkMC. (A) Quantitative determination of ATP between NTC and TFC treated hESC-SkMC. N = 8 for each condition. (B) Mitotracker signal in NTC and TFC treated hESC-SkMC. (C-F) Quantification of Mitotracker measurement between NTC and TFC for Mitotracker signal intensity (C), Mitochondria area, determined by Mitotracker area (D), Number of nuclei (E) and Normalised mitochondria area per nuclei (F). One representative biological replicate with N = 10 technical replicates is shown for each condition. (G) Representative image of NTC and TFC treated hESC-SkMC co-stained with human mitochondria antibody and MF20. (H-K) Quantification of mitochondria measurement between NTC and TFC for Mitochondria signal intensity (H), Mitochondria area within MF20+ myotubes (I), Number of myotubes (J) and Normalised mitochondria area per myotube (K). N = 3 independent biological replicates for each condition. Analysis performed with two tailed t-test. *P < 0.05, **P < 0.01, ***P< 0.001.

Nonetheless, mitochondria in TFC treated hESC-SkMC appear more elongated compared to NTC population which contains more circular mitochondria (Fig. 5B). While we did not detect any major difference in Mitotracker signal intensity between NTC and TFC (Fig. 5C), TFC treated hESC-SkMC have greater mitochondrial area than NTC (Fig. 5D), despite a similar number of nuclei analyzed (Fig. 5E), with TFC treated hESC-SkMC overall showing higher mitochondria area per nuclei compared to NTC (Fig. 5F). Similar results were obtained with a lower dose of Mitotracker (50 nM) (Supp. 7A-D).

To further confirm the results of Mitotracker measurement, we assessed mitochondrial content in myotubes by probing hESC-SkMC with human mitochondria antibody together with MF20 (Fig. 5G). TFC-treated myotubes showed greater mitochondria area compared to NTC, but no difference in mitochondria signal intensity (Fig. 5H-I). We normalized the size of mitochondria area to the number of myotubes and demonstrate TFC treated hESC-SkMC have higher mitochondria area per myotube compared to NTC (Fig. 5J-K, Supp. 7E).

Collectively, our data demonstrate that TFC treatment enhanced terminal differentiation and energy metabolism of hESC-SkMC. We describe here an improved skeletal muscle differentiation protocol leading to thicker muscle fibers associated with increased mitochondria content.

## Discussion

Muscle is a tissue with unique properties, capable of incorporating new nuclei into an already existing fiber in order to maintain its homeostasis. During human skeletal muscle generation and repair, proliferating myoblasts elongate and fuse with neighboring myoblasts forming multinucleated and contractile myotubes[49]. This phenomenon requires a constant supply of new muscle cells, and therefore the maintenance of an effective differentiation dynamic that is tightly regulated by growth factors and cytokines. Some of these factors, commonly termed myokines, are secreted by skeletal muscle (and also surrounding cells) and their role in regulating muscle mass and strength has been extensively documented[39,50]. To date, over 600 myokines have been identified[39]. In this study, we selected several myokines (VEGF, IL4, IL6 and BDNF) with reported beneficial effect on skeletal muscle and tested their effect on hESC-SkMC with or without the addition of the anabolic factors testosterone and follistatin. We report here that when combined together, testosterone, follistatin and the cocktail of myokines enhanced hESC-SkMC fusion and terminal differentiation.

We show that treatment with TFC enhanced expression of several skeletal markers in hESC-SkMC. Interestingly, multi-omics analysis revealed that TFC-treated hESC-SkMC produced distinct cellular profile compared to the individually treated hESC-SkMC. These results suggest some of these DEGs may be regulated by multiple transcription factors, each being potentially activated by one of the different treatments (T, F or C). It is possible the enhanced terminal differentiation and fusion capacity via TFC treatment is due to its ability to consistently promote expression of these DEGs, which may not be possible by a single factor alone.

Importantly, while some muscle genes were only slightly up-regulated at the mRNA level following TFC treatment, proteomics analysis showed these genes were significantly up-regulated at the protein level, suggesting that the effect of TFC may occur through translational or post translational regulation. This observation is in accordance with numerous studies showing that skeletal muscle development and function are associated with high levels of translational regulation[51,52,47].

TFC treated SkMC also exhibit higher number of nuclei per fiber and greater fusion index, suggesting these cells have better fusion capacity. This effect could at least partially be attributed to the myokine IL4 which has been shown to promote muscle growth by recruiting myoblasts and enhancing fusion[53]. TFC treated SkMC also show higher total number of nuclei compared to NTC. However, cell cycle genes and proteins are not up-regulated in these cells following TFC treatment and we did not observe enhanced proliferation in TFC-treated cells, suggesting that TFC does not induce cell proliferation. The higher number of nuclei is most likely due to a decrease in cell death during differentiation/fusion process, reflecting the protective effect of myokines on skeletal muscle. In addition to increasing muscle capillarity by its angiogenic properties[54], VEGF also exerts a direct effect on skeletal muscle and has been reported to promote the growth of myogenic fibers and protects myogenic cells from apoptosis[55]. Most myokines have an anabolic effect and prevent muscle atrophy through a variety of biological processes. IL6 is a major myokine which, following exercise, is released by skeletal muscle to maintain its homeostasis[56]. IL6 plays a central role in skeletal muscle regeneration and hypertrophy by regulating satellite cell differentiation[57]. Similarly, muscle-secreted BDNF is essential to satellite cells activation and differentiation in response to muscle injury[58]. In this study, we also added creatine, a compound found primarily in skeletal muscle that can be produced endogenously or obtained through food consumption which provides rapid energy generation during skeletal muscle contraction and is known to induce skeletal muscle hypertrophy[59,60]. The combined positive effect of these molecules likely contributed towards better differentiation of treated SkMC in this study. Surprisingly, we did not detect a significant effect of testosterone or follistatin on hESC-SkMC morphology or skeletal marker expression in the absence of the myokines cocktail, highlighting an essential role of these molecules. However, we still noticed a trend towards increased MyHC+ area in T- and F-treated hESC-SkMC. Furthermore, both compounds are known to enhance energy metabolism in skeletal muscle[61,62]. To date, over 600 myokines have been reported making it impractical to test all of them in our system in this current study. Further studies should assess the effect of other myokines on hESC differentiation into SkMC. Similarly, is well-known optimal concentration of compounds vary greatly between hPSC lines[63]. Future studies could also evaluate different concentrations of T, F and C to optimize the effect of TFC in different hPSC cell lines.

Muscle contraction is mediated by the motor protein myosin, which binds to actin and drives filament sliding[64,65]. Different types of fibers exist in skeletal muscle and they are characterized by the expression of specific MyHC ATPase isoforms. Generally, slow-twitch myofibers express the slow type I myosin ATPase and rely on oxidative metabolism. They have slow rate of contraction and are resistant to fatigue. In contrast, fast-twitch myofibers contract and fatigue rapidly, express fast type II MyHC and rely on glycolytic metabolism[66]. We previously showed by analyzing specific marker expression that our skeletal muscle differentiation protocol leads to the generation of both slow-(TNNC1+ and TNNT1+) and fast-twitch (TNNI2+, TNNT3+, TNNC2+) myofibers from hPSC[6]. In this new study, TFC treatment enhanced both slow and fast-twitch sarcomeric proteins level, which correlates with the metabolomics analysis showing that both fatty acid oxidation and glycolysis were increased. We also observed an increase in mitochondria area in TFC-treated myotubes as a possible mechanism for their enhanced energy metabolism. Since proteomics data showed oxidative phosphorylation is the top up-regulated pathway in TFC-treated SkMC, this suggests TFC may promote slow-twitch over fast-twitch myofibers due to enhanced oxidative phosphorylation capacity.

Despite enhanced terminal differentiation following TFC treatment, these cells remained immature as indicated by the presence of embryonic (MYH3) and neonatal (MYH8) markers and lack of adult myosin isoforms (MYH1 and MYH2) expression. To date, maturation of hPSC-SkMC to an adult stage can only be achieved to some extent in 3D culture environment[10,7]. Current 3D organoid cultures of hPSC-SkMC require extensive optimization of culture conditions to obtain uniform structures and may not be suitable for certain applications[67]. Future work will be focused on identifying factors that may lead to the generation of more mature, adult myotubes from hPSC.

In human MyHC level decrease with age[1] and reduced level of MyHC subsequently leads to smaller muscle mass and weakened muscular function[68,69]. The identification of factors that enhance MyHC expression or myotube density in human skeletal muscle cells would in theory reinforce muscle contractile ability and delay the progression of muscle weakness. In this study, we report an improved protocol of our skeletal muscle differentiation from hPSC, based on increasing MyHC expression in hESC-SkMC. This protocol is robust and can be employed in large scale for drug screening applications, which may assist with future studies investigating human skeletal muscle development and identification of new therapies.

## Abbreviations

hESC: human embryonic stem cell
SkMC: skeletal muscle cells
T: testosterone
F: follistatin
C: cocktail of myokines
TFC: testosterone follistatin and cocktail of myokines
MyHC: myosin heavy chain
hPSC: human pluripotent stem cells
hiPSC: human induced pluripotent stem cells
MRF: myogenic regulatory factors
IL4: interleukin-4
IL6: interleukin-6
BDNF: brain derived neurotrophic factor
VEGF: vascular endothelial growth factor
NTC: non-treated cells

## Acknowledgments

The authors thank the facilities of Sydney Microscopy and Microanalysis, Sydney Cytometry and Sydney Mass Spectrometry at the University of Sydney for their technical support. The authors would like to thank Dr. John Manion, Dr. Daniel Hesselson and Dr. Greg Neely in particular, for their assistance with overall experimental design and discussion. Finally, the authors thank the University of Sydney and the generosity of John Chong and Anne Chong for their financial support. LC is supported by a Dr. John and Anne Chong Fellowship.

## Author Contributions

TR and LC: experimental design, data collection, analysis and writing manuscript. DH and ML: mass spectrometry and data analysis. KY: metabolomics experiment and analysis. LL: calcium imaging. LC: conception, direction and writing manuscript.

---

Table S1. Table of Antibodies Used

Table S2. Table of Growth Factors Used for Skeletal Muscle Differentiation

Table S3. Primer Sequences for RT-qPCR

