## Supplemental material for "Anabolic Factors and Myokines Improve Differentiation of Human Embryonic Stem Cell Derived Skeletal Muscle Cells"

#### Supplemental. 1

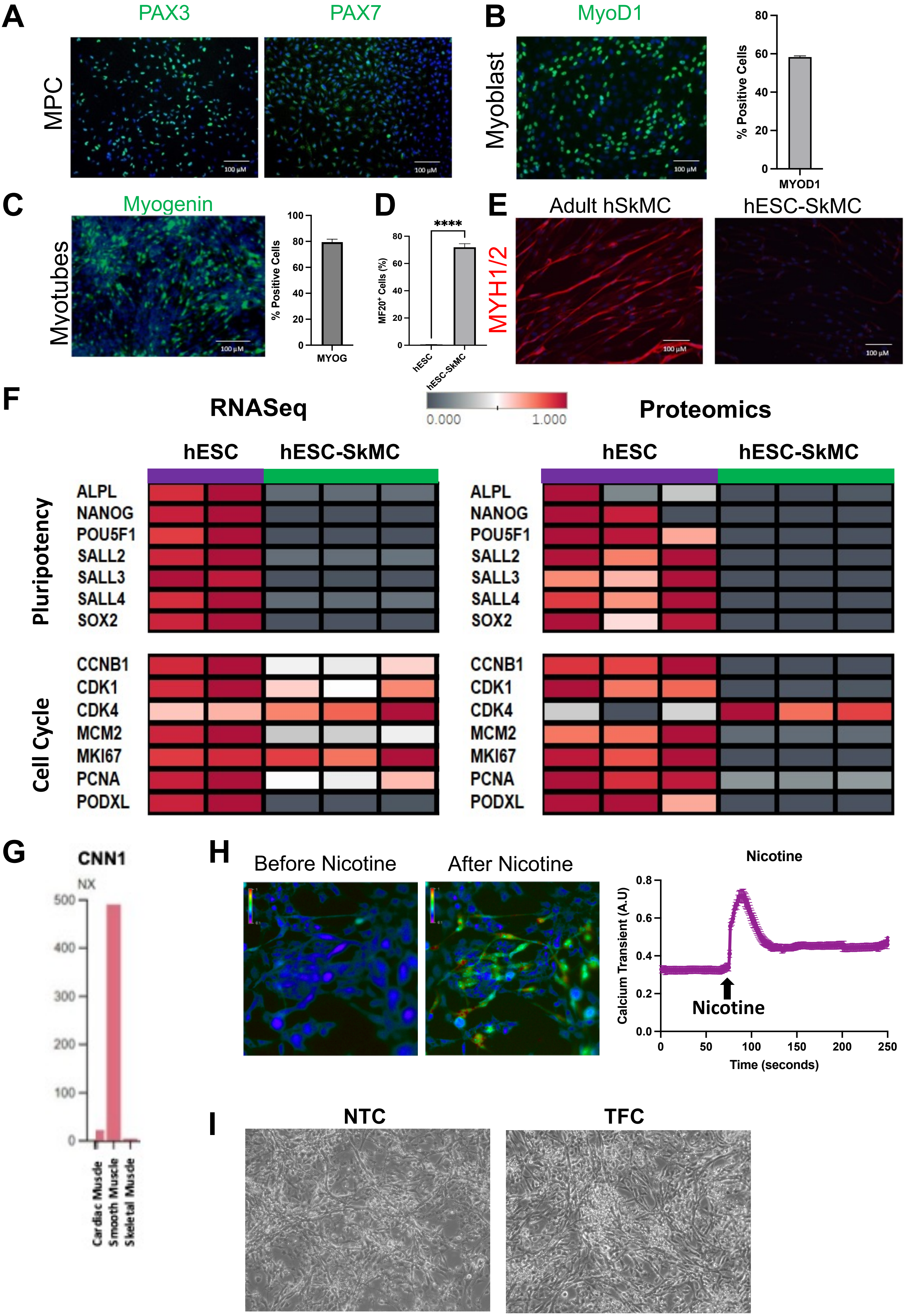

Supplemental. 2

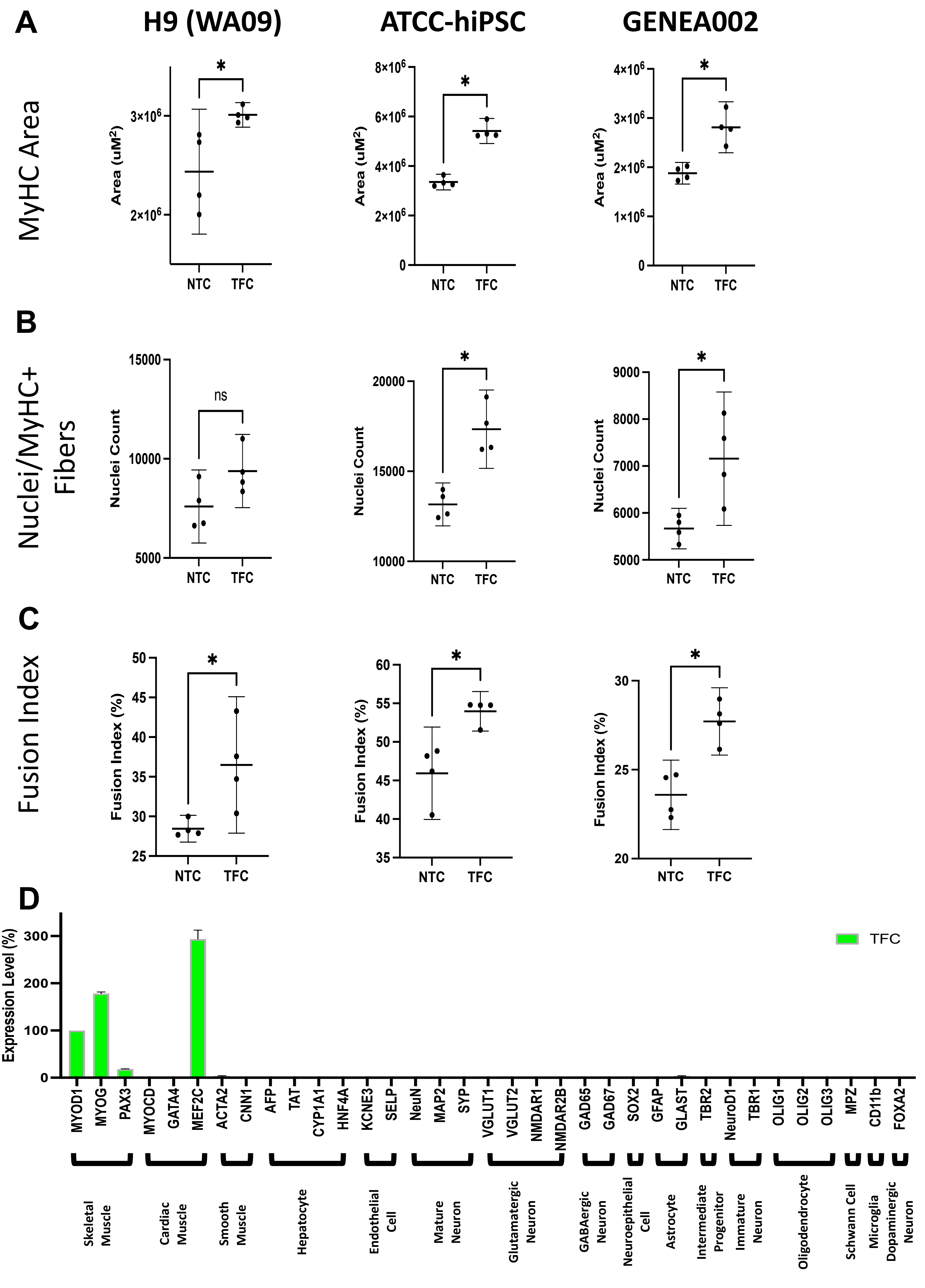

Supplemental. 3

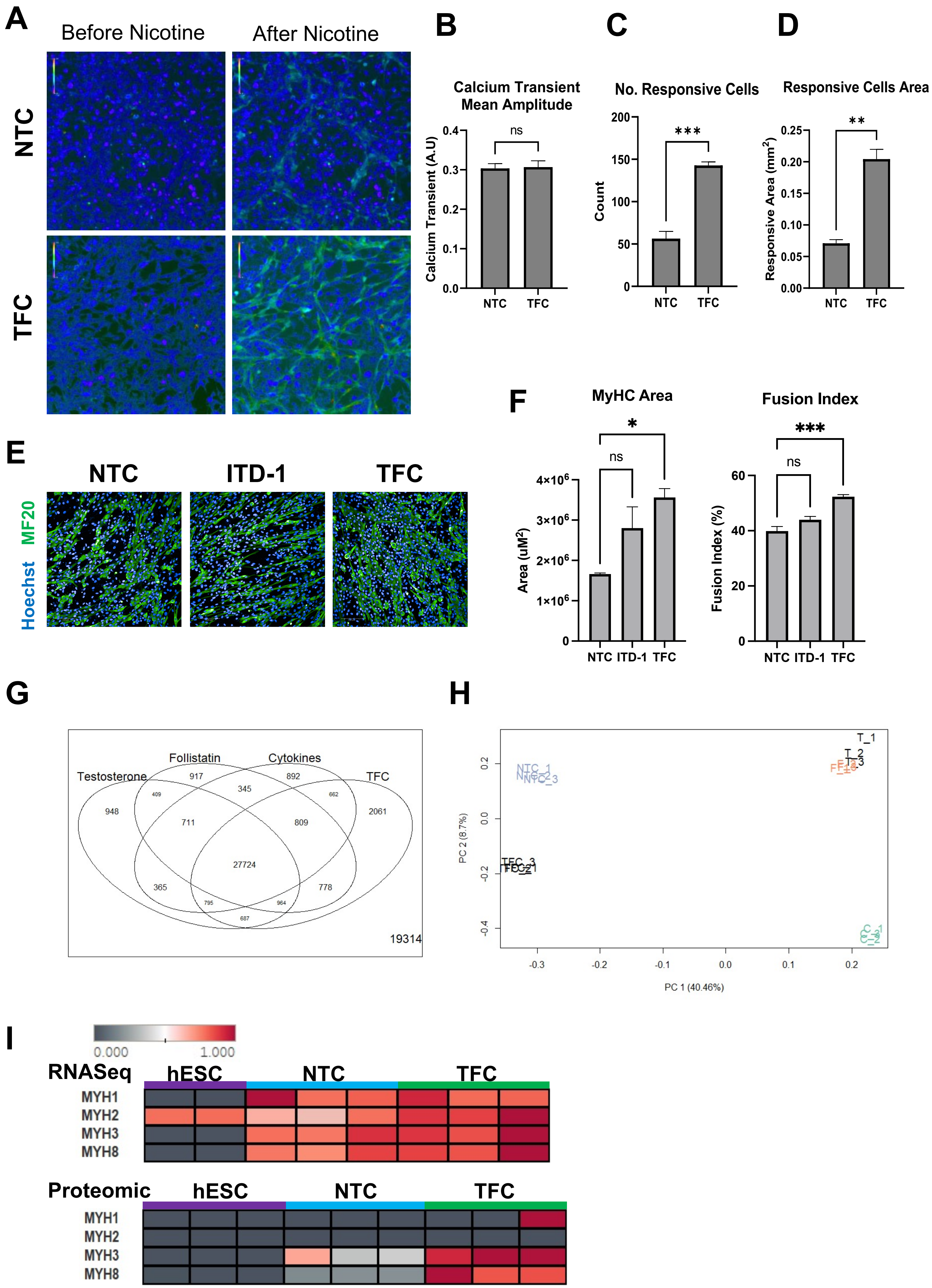

### Supplemental. 4

**A**

| Condition | No. Proteins not detected in NTC |
|---|---|
| Testosterone | 195 |
| Follistatin | 260 |
| Cocktail of myokines | 214 |
| T and F | 105 |
| T and C | 98 |
| F and C | 125 |
| T and F and C | 80 |

**B**

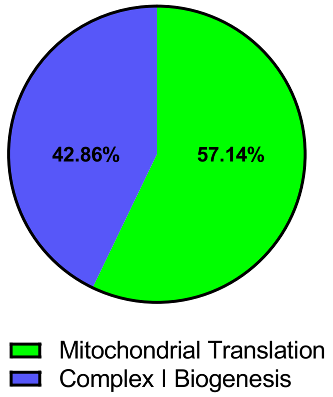


**C**

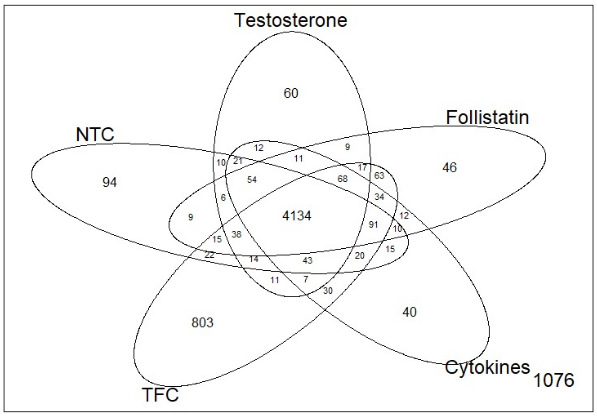


**D**

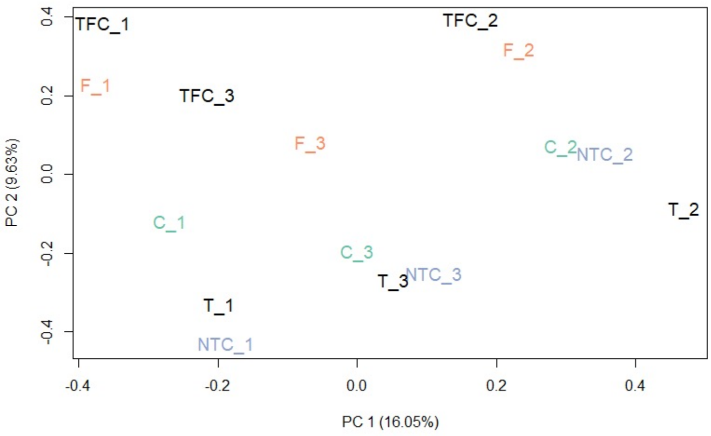


**E**

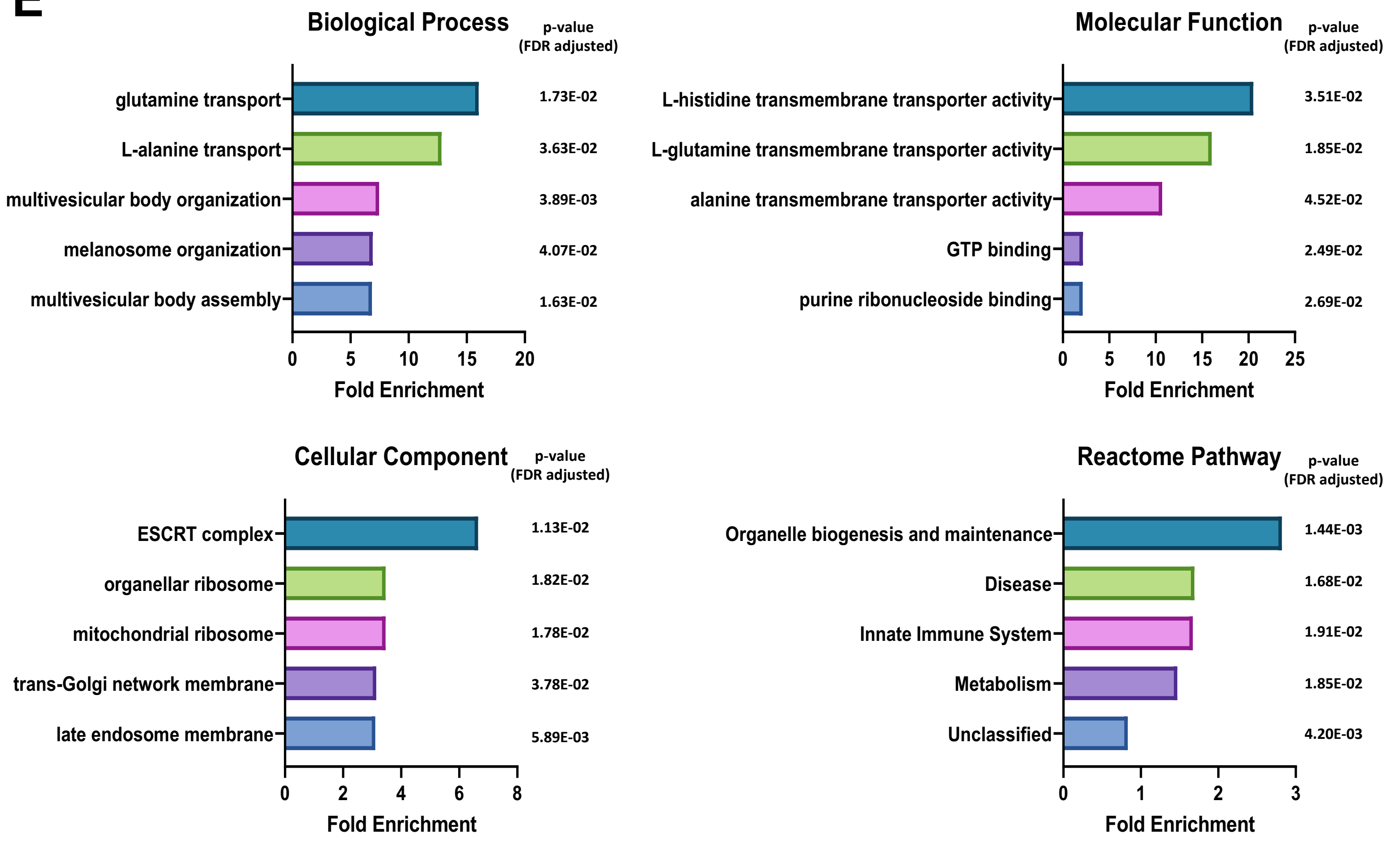

**Supplemental. 4**

**F**

### TFC v NTC

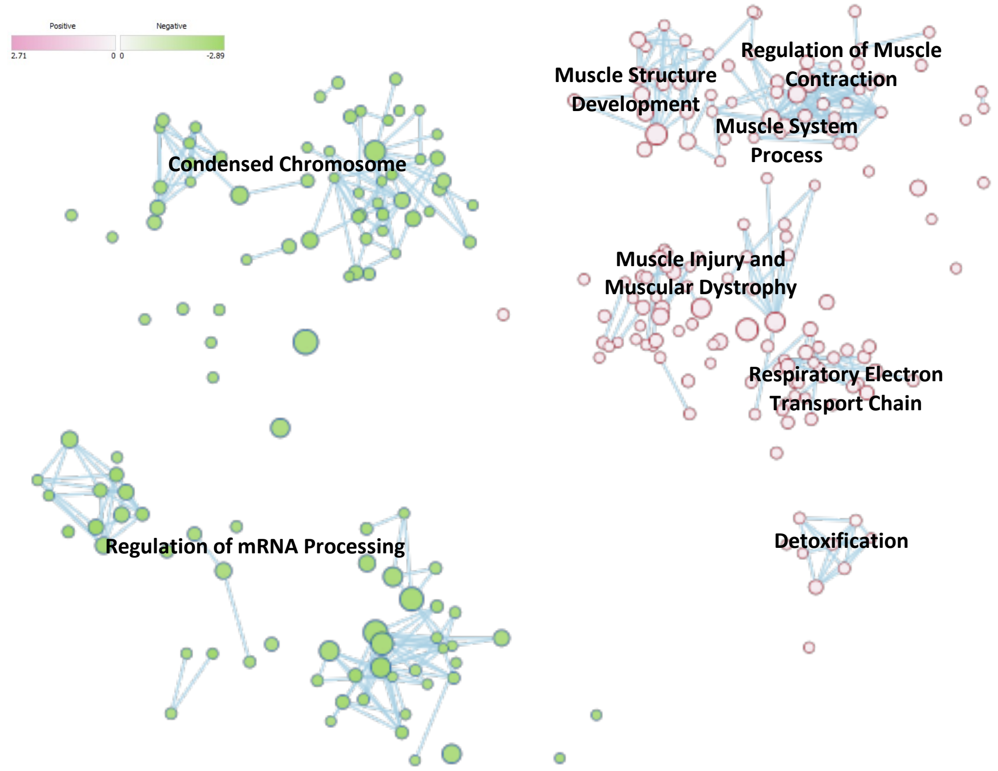


**G**

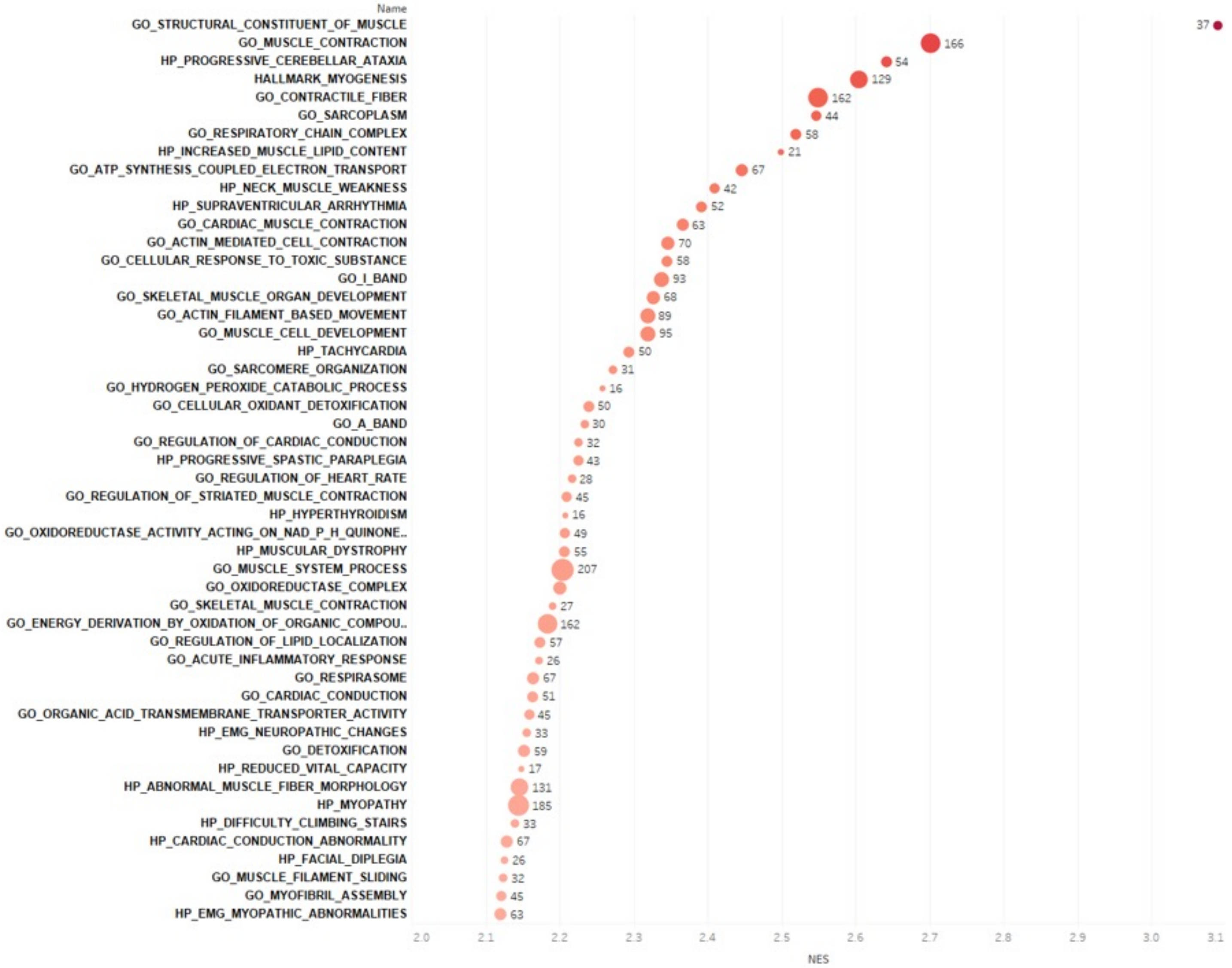

**Supplemental. 5**

A

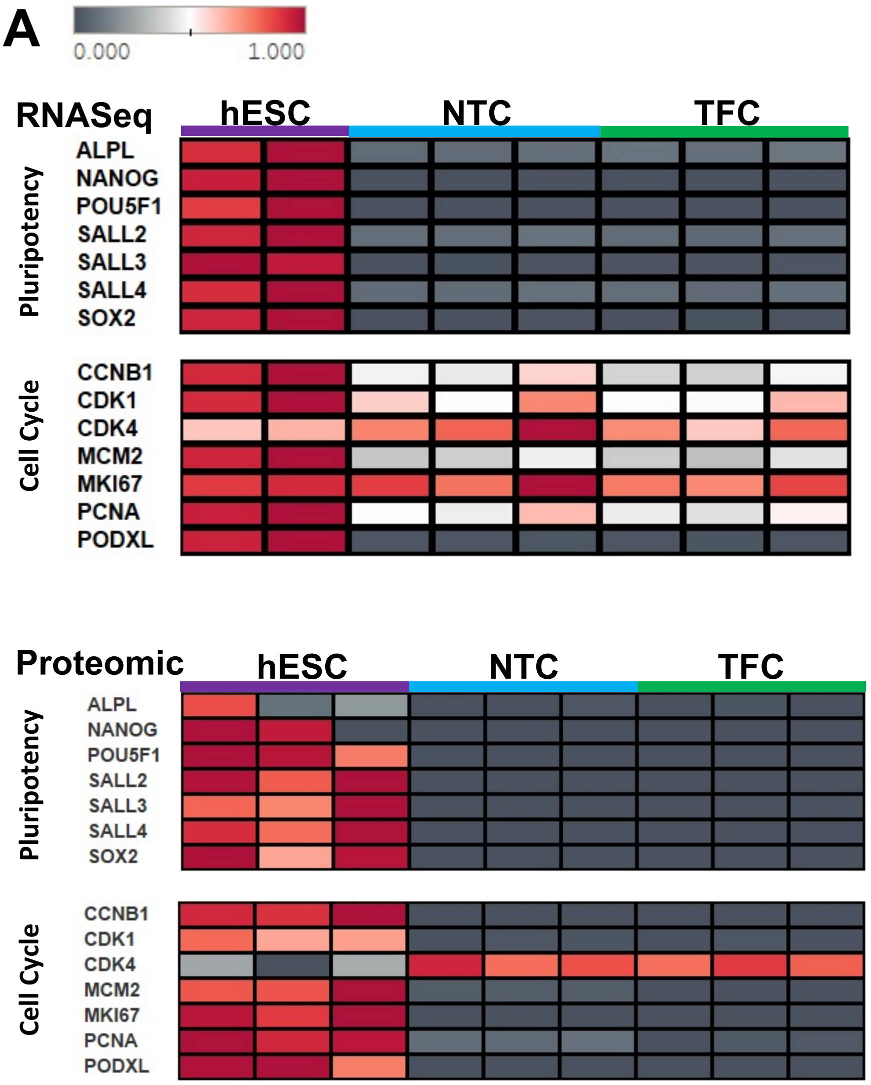


B

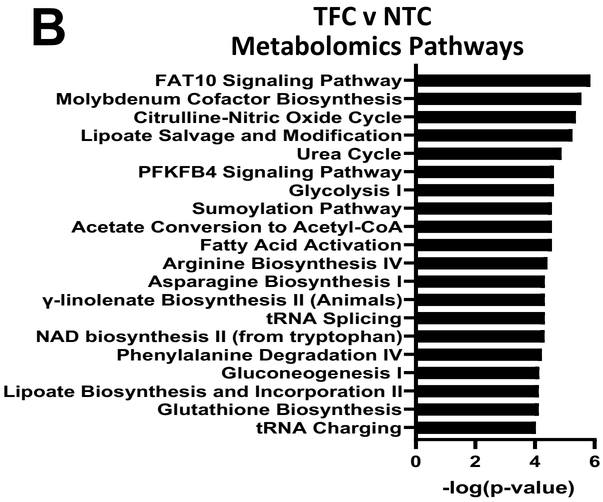

#### Supplemental. 6

**A**

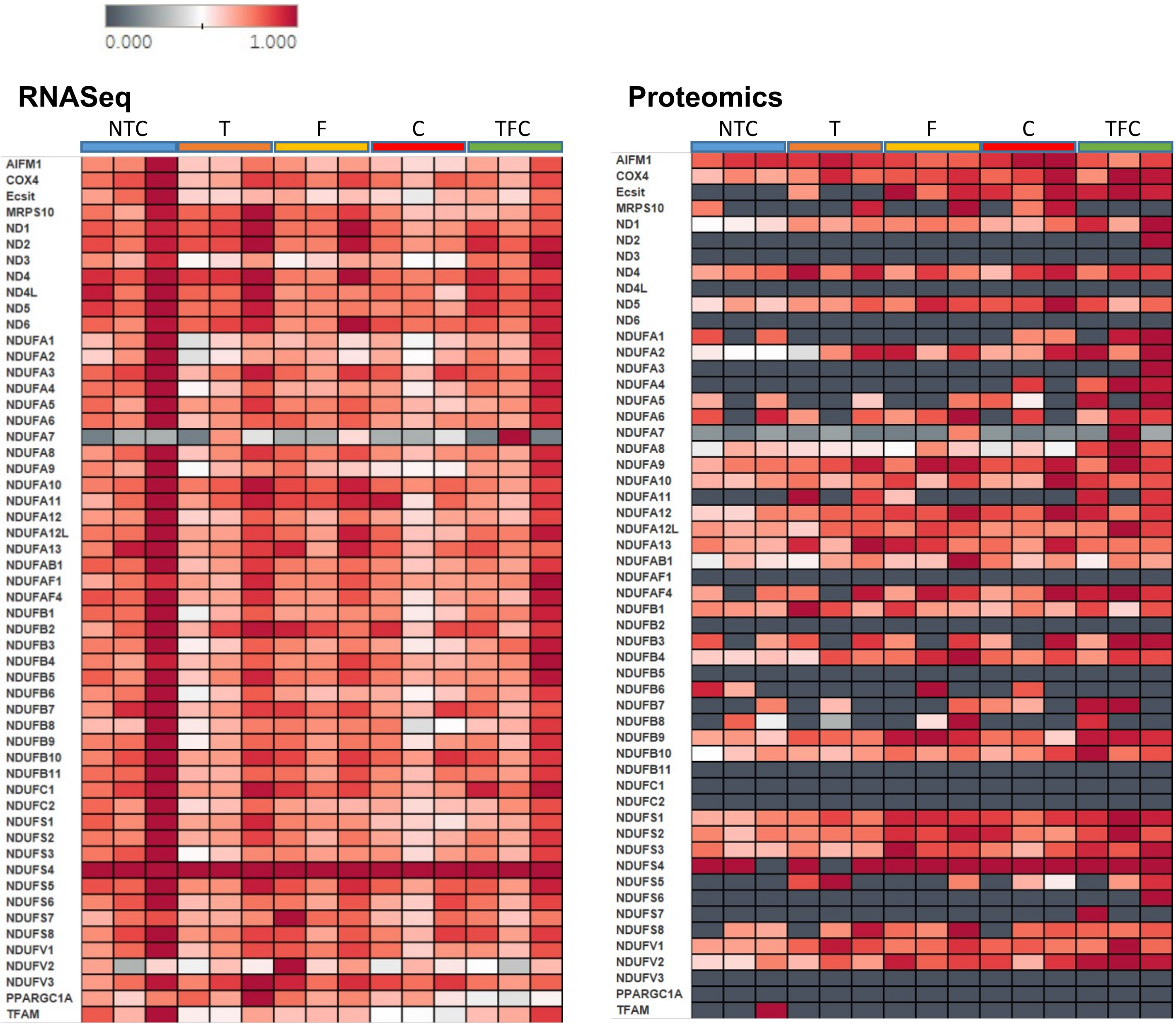


**B**

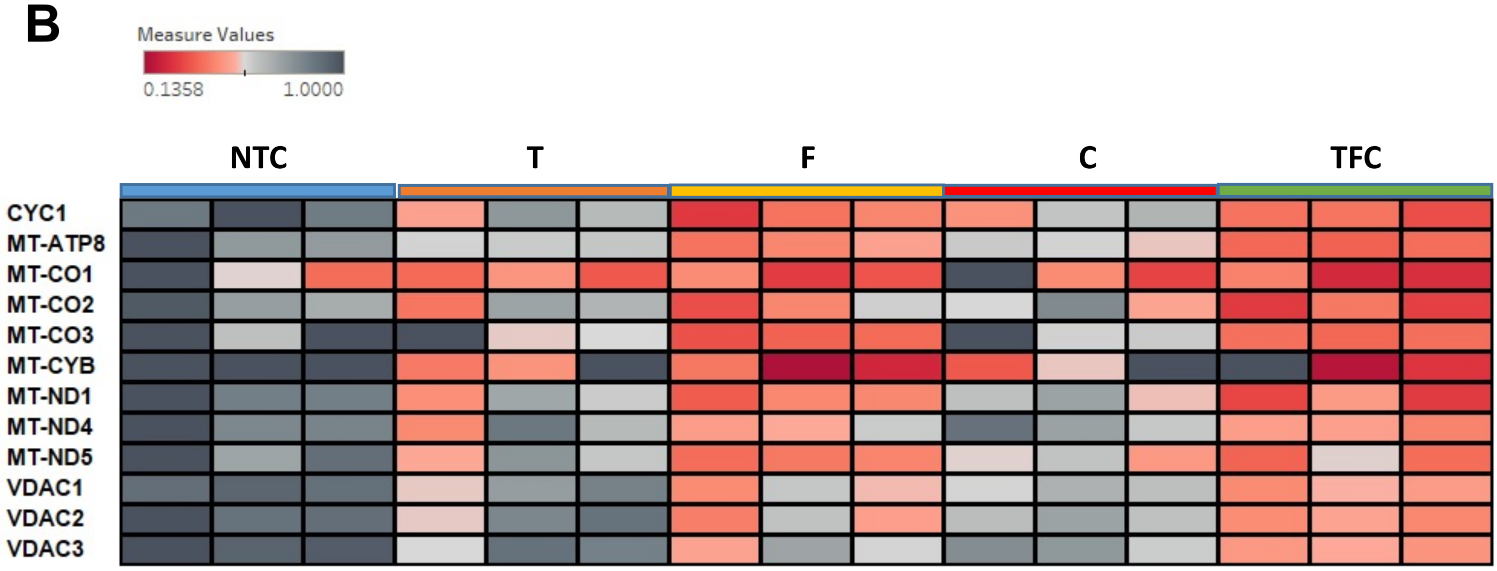

#### Supplemental. 7

**A** Mitotracker DeepRed staining (entire well)

NTC TFC

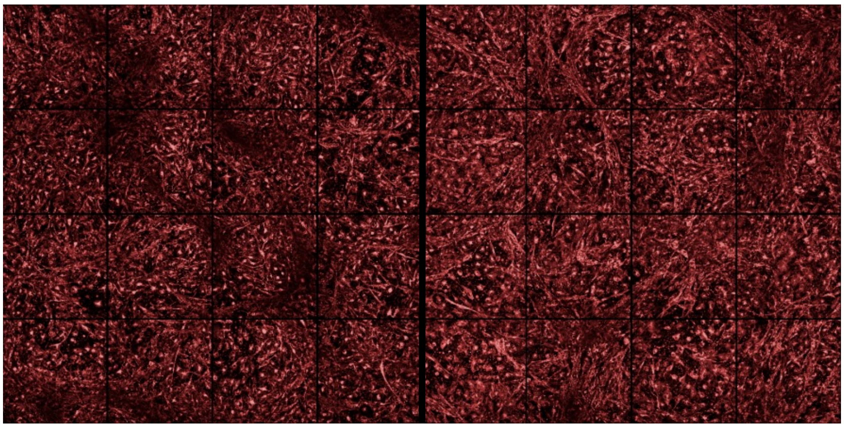

Mitotracker DeepRed (50nM)

**B** **Mitochondria area**

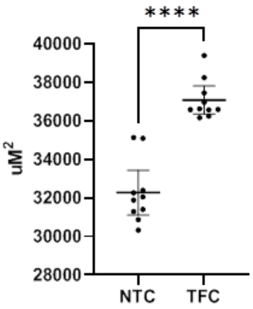


**C** **Number of Nuclei**

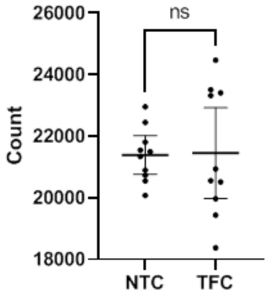


**D** **Mitochondria area per Nuclei**

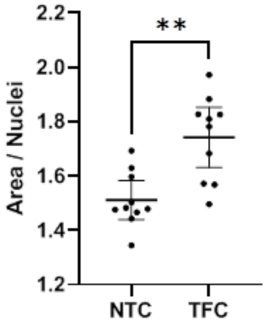


**E** Human Mitochondria Ab staining (entire well)

**Hoechst** **MF20** **Human Mitochondria**

**NTC**

**TFC**

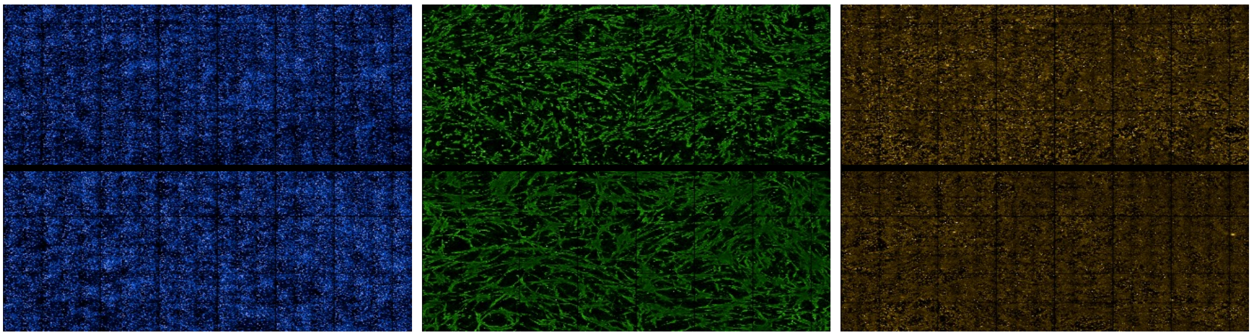

**Merged**

**NTC**

**TFC**

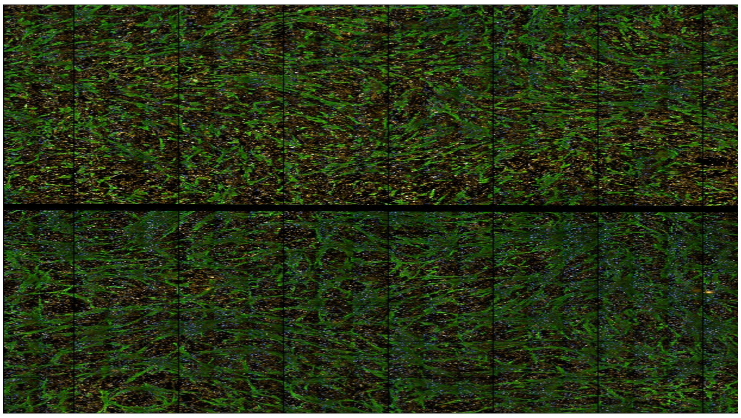

#### Table S1. Table of Antibodies Used

| Name | Cat# | Supplier | Concentration | Application |
|---|---|---|---|---|
| Dystrophin | MANDRA1(7A10) | DSHB | 1:50 | IF |
| a-Actinin | A7811 | Sigma | 1:800 | IF |
| MF20 | MF20 | DSHB | 1:400 | IF |
| MYH3 | F1.652 | DSHB | 1:50 | IF |
| MYH8 | N3.36 | DSHB | 1:50 | IF |
| PAX3 | MAB2457 | R&D Systems | 1:100 | IF |
| PAX7 | MAB1675 | R&D Systems | 1:100 | IF |
| MyoD1 | 554130 | BD Pharmingen | 1:200 | IF |
| Desmin | 550626 | BD Pharmingen | 1:200 | IF |
| Myogenin | 556358 | BD Pharmingen | 1:200 | IF |
| MyH1 | 6H1 | DSHB | 1:200 | IF |
| Human Mitochondria | ab92824 | Abcam | 1:800 | IF |
| Mitotracker Deep Red | M22426 | ThermoFisher | 50nM & 250nM | IF |
| Donkey anti-Mouse IgG | A21202 | ThermoFisher | 1:500 | IF |
| Goat anti-Mouse IgG2b | A21242 | ThermoFisher | 1:500 | IF |

#### Table S2. Table of Growth Factors Used

| Phase 1 | Cat# | Supplier | Concentration |
|---|---|---|---|
| Skeletal Muscle Basal Media | C-23260 | PromoCell | |
| Horse Serum | 16050122 | ThermoFisher | 5% |
| CHIR99021 | 72052 | StemCell Technologies | 3uM |
| ALK5 Inhibitor | 73792 | StemCell Technologies | 2uM |
| Human Recombinant Epidermal Growth Factor | 130-097-749 | Miltenyi Biotec | 10ng/mL |
| Insulin | I9278 | Sigma | 10ug/mL |
| Dexamethasone | D4902 | Sigma | 0.4ug/mL |
| Ascorbic Acid | A5960 | Sigma | 200uM |
| **Phase 2** | **Cat#** | **Supplier** | **Concentration** |
| Skeletal Muscle Basal Media | C-23260 | PromoCell | |
| Horse Serum | 16050122 | ThermoFisher | 5% |
| Insulin | I9278 | Sigma | 10ug/mL |
| Human Recombinant Epidermal Growth Factor | 130-097-749 | Miltenyi Biotec | 10ng/mL |
| Human Recombinant Hepatocyte Growth Factor | 100-39H | PeproTech | 20ng/mL |
| Human Recombinant Platelet Derived Growth Factor | 100-00AB | PeproTech | 10ng/mL |
| Human Recombinant Basic Fibroblast Growth Factor | 100-18B | PeproTech | 20ng/mL |
| Oncostatin | 130-114-939 | Miltenyi Biotec | 20ng/mL |
| Insulin-like Growth Factor 1 | 130-093-886 | Miltenyi Biotec | 10ng/mL |
| SB431542 | 72232 | StemCell Technologies | 2uM |
| Ascorbic Acid | A5960 | Sigma | 200uM |

#### Table S2. Table of Growth Factors Used

| Phase 3 | Cat# | Supplier | Concentration |
|---|---|---|---|
| Skeletal Muscle Basal Media | C-23260 | PromoCell | |
| Insulin | I9278 | Sigma | 10ug/mL |
| Oncostatin | 130-114-939 | Miltenyi Biotec | 20ng/mL |
| Necrosulfonamide | 5025/10 | Torcis | 50nM |
| Ascorbic Acid | A5960 | Sigma | 200uM |

| TFC | Cat# | Supplier | Concentration |
|---|---|---|---|
| Testosterone | T1500 | Sigma | 400ng/mL |
| Follistatin | SRP3045 | Sigma | 300ng/mL |
| Creatine | C0780 | Sigma | 1mM |
| Interleukin-6 | 200-06 | PeproTech | 150ng/mL |
| Interleukin-4 | 200-04 | PeproTech | 20ng/mL |
| Brain Derived Neurotrophic Factor | 450-02 | PeproTech | 20ng/mL |
| Vascular Endothelial Growth Factor | 100-20 | PeproTech | 25ng/mL |

**Table S3. Primer Sequences**

| Gene | | Sequences |
|---|---|---|
| MyoD1 | F | CGGCATGATGGACTACAGCG |
| | R | CAGGCAGTCTAGGCTCGAC |
| MYH3 | F | ATTGCTTCGTGGTGGACTCAA |
| | R | GGCCATGTCTTCGATCCTGTC |
| MYH8 | F | CCAAAACAAGCCGTTTGATGC |
| | R | AGCACTCCAGGCTCGTGTA |
| Desmin | F | CAGGAGATGATGGAATACCG |
| | R | TTCTTGGTATGGACCTCAGA |
| TNNT | F | TGCTCCCGCCAAAGATCCC |
| | R | TCTTCCGCTGCTCGAAATGTA |
| DMD | F | GGTGGGAAGAAGTAGAGGAC |
| | R | ACCCAGCTCAGGAGAATCTT |
| CLDN11 | F | CGGTGTGGCTAAGTACAGGC |
| | R | CGCAGTGTAGTAGAAACGGTTTT |
| RBP4 | F | AGGAGAACTTCGACAAGGCTC |
| | R | GAGAACTCCGCGACGATGTT |
| CA4 | F | CTGGTGCTACGAGGTTCAAGC |
| | R | GAAGAAGAAGCGTCCCAGTTT |
| PRF1 | F | GGCTGGACGTGACTCCTAAG |
| | R | CTGGGTGGAGGCGTTGAAG |
| SCGN | F | AACTGGGTACTGATGACACGG |
| | R | TCTTTAGAGGCATCTTGGGTAGT |
| ALOX15 | F | GGGCAAGGAGACAGAACTCAA |
| | R | CAGCGGTAACAAGGGAACCT |

**Table S3. Primer Sequences**

| Gene | | Sequences |
|---|---|---|
| FABP4 | F | ACTGGGCCAGGAATTTGACG |
| | R | CTCGTGGAAGTGACGCCTT |
| ST8SIA3 | F | TCGCCCTGCTGATTTTATCG |
| | R | AGCGCAAATTGTGACCGGA |
| VDR | F | GTGGACATCGGCATGATGAAG |
| | R | GGTCGTAGGTCTTATGGTGGG |
| DUOX2 | F | CTGGGTCCATCGGGCAATC |
| | R | GTCGGCGTAATTGGCTGGTA |
| NEB | F | CCCCTACATTGCACACAGTCA |
| | R | GCGTATGGCTGTCCTTTTGTT |
| MYOG | F | GGTTGCCCAGCGAATGC |
| | R | TGATGCTGTCCACGATGGA |
| GAPDH | F | TCAAGAAGGTGGTGAAGCAGG |
| | R | ACCAGGAAATGAGCTTGACAAA |

**Supplemental. 1** (A-C) Immuno-staining of cells with skeletal muscle markers during skeletal muscle differentiation. PAX3 and PAX7 (A), MyoD1 (B) and MYOG (C). (D) Immuno-staining of adult primary SkMC and hESC-SkMC with 6H1 antibody (DSHB) marking both adult MyHC isoforms MYH1 and MYH2. (E) Skeletal muscle differentiation is associated with down-regulation of pluripotency and cell cycle genes. (F) CNN1 is expressed in all three types of muscle. Data obtained from The Human Protein Atlas website https://www.proteinatlas.org/ENSG00000130176-CNN1. (G) Representative image of calcium transients following the addition of nicotine. (H) Representative brightfield image of untreated and treated hESC-SkMC. Shown is one representative experiment of at least 3 biological replicates. MYOG was performed in three technical replicates.

**Supplemental. 2** TFC enhanced MyHC expression in hPSC lines including H9 (WA09), GENEA002, and ATCC-hiPSC. (A) MyHC area. (B) Nuclei within MyHC+ fibers. (C) Fusion index. One representative biological replicate with N = 4 technical replicates is shown for each condition. Analysis performed with two tailed t-test. *P < 0.05, **P < 0.01, ***P< 0.001. (D) TFC treated hESC-SkMC express markers specific to skeletal muscle lineage. Shown is data pooled from 3 independent biological replicates.

**Supplemental. 3** (A) Representative image of calcium transients following the addition of nicotine between NTC and TFC treated hESC-SkMC. (B) No difference in calcium transient mean amplitude between NTC and TFC. (C) More cells in TFC treated hESC-SkMC responded to nicotine stimulation compared to NTC. (D) Greater cellular area in TFC treated hESC-SkMC responded to nicotine stimulation compared to NTC. N = 3 for each condition. Analysis performed with two tailed t-test. *P < 0.05, **P < 0.01, ***P< 0.001. (E) Representative image of hESC-SkMC treated with ITD-1 (5μM) or TFC compared to control. (F) Quantification of MyHC area and fusion index between control, ITD-1 and TFC treated hESC-SkMC. Statistical analysis performed using One-way ANOVA with Benjamini-Hochberg FDR correction. *P < 0.05, **P < 0.01, ***P < 0.001. (G) Venn diagram showing common and divergent genes between different treatments (T, F, C and TFC) in RNASeq. (H) Principal Component Analysis (PCA) of untreated and treated hESC-SkMC in RNASeq. (I) NTC and TFC express high level of MYH3 and MYH8 and minimal level of MYH1 and MYH2.

**Supplemental. 4** (A) Individual treatment induced expression of genes not detected in NTC in proteomics. (B) Gene Ontology analysis of the pathways associated with these 80 genes up-regulated in all three individual treatment. (C) Venn diagram showing common and divergent proteins between different treatments (T, F, C and TFC). (D) PCA of untreated and treated hESC-SkMC in proteomics. (E) Gene Ontology analysis of genes detected only in TFC treatment in proteomics. (F) GSEA analysis of major differentially regulated pathways between TFC and NTC in proteomics. (G) GSEA analysis of top enriched gene sets in phenotype TFC compared to NTC. Shown is data pooled from three 3 independent biological replicates.

**Supplemental. 5** (A) Heatmap comparison of pluripotency and cell cycle markers between hESC, NTC and TFC. (B) Top enriched metabolomics pathways between TFC and NTC (IPA Log2FC>1, padj<0.05). Shown is data pooled from three independent biological replicates.

**Supplemental. 6** (A) Heatmap showing the expression of various mitochondrial genes between treatments at the RNA and protein level. (B) Heatmap comparison of nuclear (Histone H4) vs mitochondrial protein ratio in untreated (NTC) and treated hESC-SkMC. Shown is data from three independent biological replicates.

**Supplemental. 7** Comparison of mitochondria abundance between NTC and TFC treated hESC-SkMC. (A) Full well view of a representative Mitotracker staining of NTC and TFC treated hESC-SkMC. (B-D) Quantification of mitochondria area between NTC and TFC treated hESC-SkMC using 50nM of Mitotracker Deep Red for area (B), Number of nuclei (C), and Normalised mitochondria area per nuclei (D). One representative biological replicate with N = 10 technical replicate is shown for each condition. Analysis performed with two tailed t-test. $^*P < 0.05$, $^{**}P < 0.01$, $^{***}P < 0.001$. (E) Representative image of human mitochondria antibody co-stained with MF20 in NTC and TFC treated hESC-SkMC.
